# Transmembrane domain composition reflects subcellular localization of SNARE proteins

**DOI:** 10.64898/2026.03.24.713902

**Authors:** Christian Baumann, Carlos Pulido-Quetglas, Dirk Fasshauer

**Affiliations:** Department of Computational Biology, University of Lausanne, CH-1015 Lausanne, Switzerland

## Abstract

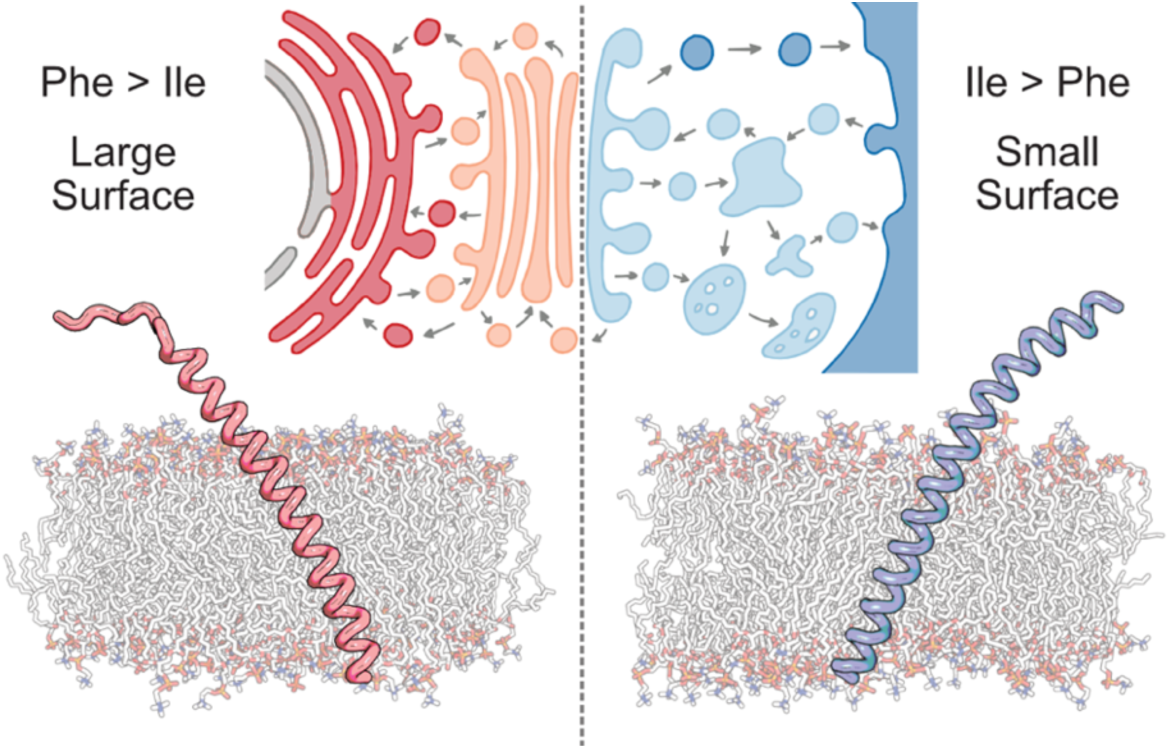

Transmembrane domains (TMDs) anchor proteins within membranes of distinct composition and biophysical properties, necessitating adaptation to each membrane’s unique environment. SNARE proteins, the core mediators of eukaryotic membrane fusion, provide an ideal system to study such adaptations: they are ubiquitous across Eukaryota, with paralogs localized to different compartments that have evolved independently to match their membrane context. Here, we combine statistical analyses of approximately 14,000 SNARE TMDs with MD simulations of 28 human SNARE TMDs to uncover how TMDs have adapted to distinct membrane environments. We identify a clear compositional dichotomy that separates SNAREs of the early secretory pathway (ER and Golgi apparatus) from those of the late secretory and endosomal pathways: early TMDs are enriched in bulky Phe residues, whereas late TMDs are dominated by smaller Ile residues. The larger Phe side chains likely enhance hydrophobic interactions in loosely packed ER and Golgi membranes, while Ile likely prevents disruption of the tight lipid-lipid packing in the plasma and endosomal membranes, thereby contributing to protein localization. TMD lengths derived from MD simulations differ from those inferred by sequence-based approaches and show limited variations across compartments. Together, these results reveal how variations in membrane composition shaped SNARE TMDs across eukaryotic compartments.

## Introduction

Compartmentalization enables eukaryotic cells to perform complex biochemical processes in distinct local environments. Despite being physically separated, cellular compartments are not isolated; they continuously communicate and exchange materials through dynamic processes. One major mode of exchange occurs through vesicles that bud from one compartment and fuse with another. Membrane fusion is a tightly regulated process mediated by specific proteins, most notably the SNARE (soluble *N*-ethylmaleimide-sensitive factor attachment protein receptor) family. About 20 proteins of this family were already present in the last eukaryotic common ancestor (LECA), and have since diversified into numerous lineage-specific variants, totaling 38 in humans (Fig. 1A).^1–4^

**Figure 1.**
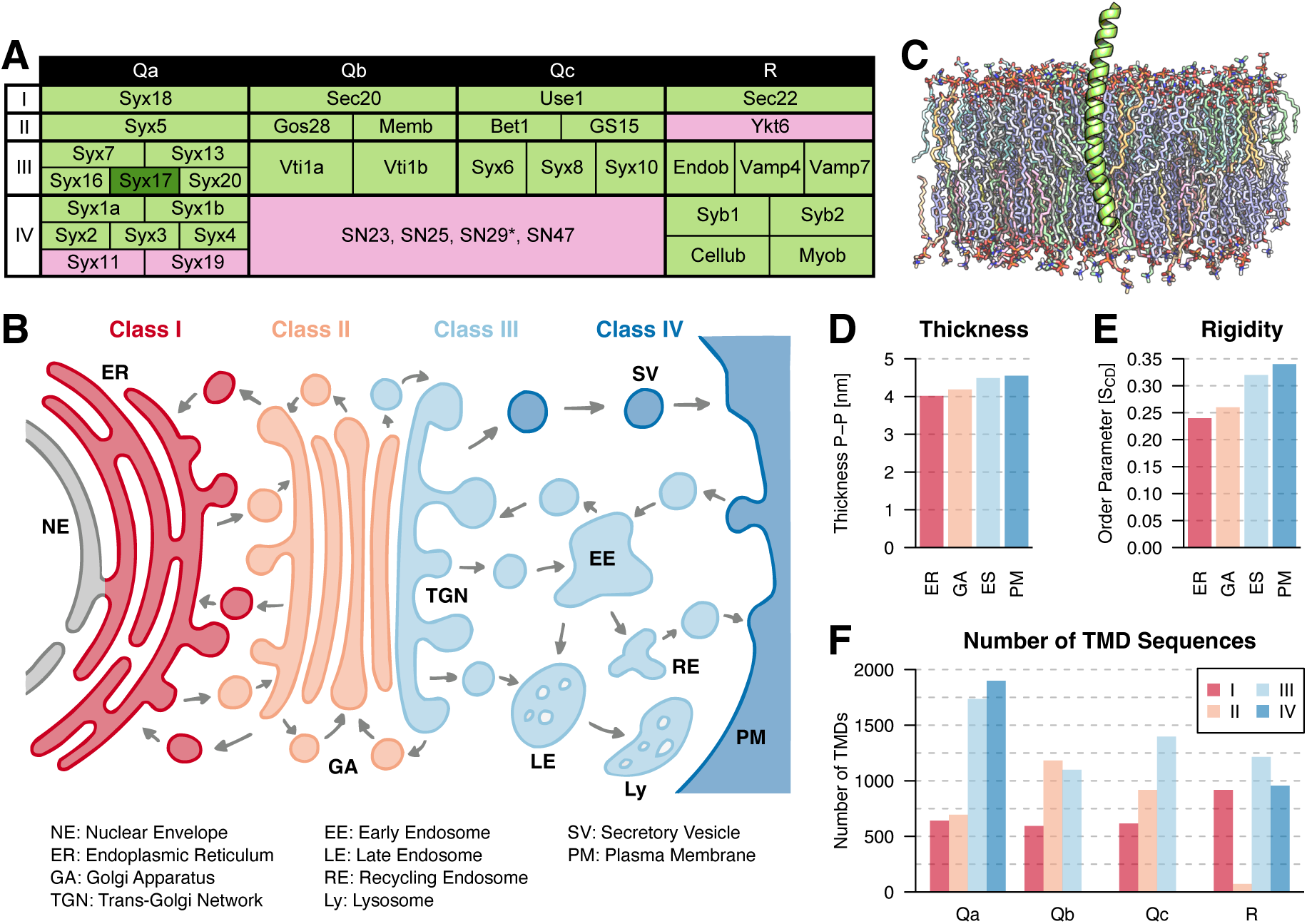
Overview of human SNAREs, membrane environments, and dataset composition. **(A)** SNAREs in humans. Proteins containing transmembrane domains (TMDs) are shown in green, whereas lipid-anchored SNAREs are shown in pink. The TMD of Syx17 consists of two α-helices rather than one. SN29 lacks any type of membrane anchor. **(B)** Schematic of cellular organization indicating the compartments in which SNAREs function: class I at the endoplasmic reticulum (ER) (red), class II at the Golgi apparatus (GA) (orange), class III at the *trans*-Golgi network (TGN) and endosomal system (ES) (light blue), and class IV at the plasma membrane (PM) and secretory vesicles (SVs) (dark blue). **(C)** *C*-terminal region of Syb2 embedded in a membrane, shown as an example of a SNARE TMD. The portion of the α-helix buried within the lipid bilayer is referred to as the TMD. **(D)** Membrane thickness in mammalian model membranes, measured as the distance between the phosphate atoms of opposing lipid leaflets. Based on data from Pogozheva *et al*.^27^ **(E)** Lipid rigidity in mammalian model membranes, represented by the deuterium order parameter; smaller values indicate loosely packed, more flexible lipids. Based on data from Pogozheva *et al*.^27^ **(F)** Number of SNARE TMD sequences analyzed in this study, categorized by type (Qa, Qb, Qc, R) and class (I, II, III, IV).

SNAREs mediate membrane fusion by forming tetrameric complexes composed of SNAREs anchored in both participating membranes.^5,6^ The assembly of these complexes in a zipper-like fashion draws the two membranes into close proximity, ultimately resulting in their fusion. Each tetramer consists of four distinct SNARE types (Qa, Qb, Qc, and R) that interact in a defined manner.^7,8^ The four types can be distinguished by characteristic sequence signatures. Their names stem from the conserved amino acids at the central layer of the SNARE complex: Q-SNAREs contain a Gln, whereas R-SNAREs carry an Arg residue. The four SNARE types can each be divided into four classes, which can be assigned to specific intracellular localizations (Fig. 1B): class I at the endoplasmic reticulum (ER), class II at the *cis*- and medial Golgi apparatus (GA), class III at the *trans*-Golgi network (TGN) and endosomal system (ES), and class IV at the plasma membrane (PM) and secretory vesicles (SVs).^1^

Most SNAREs are anchored in membranes via a *C*-terminal transmembrane domain (TMD) (Fig. 1C). Typically, the SNARE TMD consists of a single α-helix that spans the lipid bilayer. Notable exceptions include Syx17, which contains a two-helix TMD, and several SNAREs that are anchored via lipidation of cysteine residues.^3,9^ Like other tail-anchored proteins, SNAREs are targeted post-translationally to the ER membrane via the GET pathway and reach their destinations within the secretory or endocytic pathway through membrane trafficking.^10,11^ SNARE-mediated membrane fusion is highly specific in that vesicles fuse only with defined target compartments, requiring the correct spatial localization of SNAREs within the cell. Two distinct mechanisms ensure correct SNARE sorting. First, specific sequence or structural motifs are recognized by, for example, sorting receptors or coat proteins.^12,13^ One example is the KDEL motif, which promotes ER retention and is found in a subset of Sec20 proteins.^14,15^ Second, the physicochemical properties of the TMD itself can drive lateral segregation within membranes, enabling sorting and, in some cases, being sufficient for correct SNARE localization.^16–19^

Eukaryotic endomembranes are complex mixtures composed of hundreds to thousands of distinct lipid species.^20^ Lipid compositions vary between species, tissues, organelles, leaflets, and even membrane subdomains, with cells devoting substantial resources to maintain specific lipid distributions.^20,21^

Despite this diversity, membranes can be broadly categorized into two regimes with distinct characteristics and biological roles.^22,23^ The first regime comprises loosely packed, thin membranes enriched in unsaturated lipids, typical of early secretory compartments such as the ER and GA. The second regime includes tightly packed, thick membranes enriched in saturated lipids, sphingolipids, sterols, and negatively charged lipids, particularly in the cytosolic leaflet. This membrane type characterizes the PM, TGN, and endosomal membranes, with the transition between the two regimes occurring from the medial Golgi stacks to the TGN. Membrane composition is not uniform across these two regimes but instead follows a continuous gradient along the secretory route from the ER to the PM: lipid monounsaturation decreases, while lipid saturation, sterol and sphingolipid content, transbilayer lipid asymmetry, and negative surface charge on the cytoplasmic leaflet all increase.^24–26^ These compositional gradients produce distinct physicochemical effects, including progressive increases in membrane thickness (Fig. 1D) and lipid rigidity (Fig. 1E) along the secretory pathway.^27^

Membrane proteins evolved to match the physical and chemical properties of their native membranes.^28–31^ Research over the past decades has revealed several key principles, including that TMD length increases along the secretory pathway and contributes to protein sorting, that positively charged residues are predominantly found at the cytosolic side of the PM, and that TMD volumes are asymmetrically distributed between the two leaflets of the PM.^28,32–35^ These studies did not focus specifically on SNAREs; however, Sharpe *et al.* showed that SNARE TMDs follow similar trends to those observed for other bitopic membrane proteins.^28^ SNAREs provide an excellent model system for examining how membranes shape protein properties for two reasons: (1) SNARE classes do not form monophyletic groups, and thus highlight convergent adaptations to their native membranes.^1,36^ (2) All SNAREs perform the same conserved role in membrane fusion, minimizing the influence of functional divergence on potential differences in their TMDs. Moreover, we have established an extensive collection of classified SNARE sequences across Eukaryota, which allows for robust comparative analyses of their TMD sequences.^1,37^

In this study, we analyze SNARE TMDs in terms of their amino acid composition, TMD length and lipid-accessible surface area (LASA). Our results reveal that TMD amino acid composition and LASA reflect the two membrane regimes: ER and GA SNAREs are enriched in Phe and have a large LASA, whereas ES and PM SNAREs show higher levels of Ile and a small LASA, which likely reflects whether strong or weak protein-lipid interactions are advantageous in their native membrane environment. TMD lengths are found to be surprisingly similar across classes, and short TMDs are not restricted to the early secretory pathway.

## Results

### Amino acid composition

SNAREs mediate membrane fusion between specific vesicle-compartment pairs and therefore must be localized at the correct membranes within the cell. Since cellular membranes differ in their biophysical properties, we wondered whether these differences are reflected in the amino acid composition of SNARE TMDs. To address this question, we analyzed the amino acid composition of 13,941 SNARE TMD sequences from our extensive in-house SNARE collection with respect to their site of action within the cell (Fig. 1F). This analysis reveals pronounced class-specific differences: classes I and II, that is, SNAREs of the ER and GA, contain a higher proportion of Phe, whereas classes III and IV, that is, SNAREs of the TGN, ES, SVs and PM, are enriched in Ile (Fig. 2A), although considerable variation in individual amino acid frequency exists among TMDs within each class, resulting in substantial overlap between classes. Our results closely align with an earlier study by Munro, who analyzed amino acid composition of GA and PM TMDs using a much smaller dataset.^38^ Particularly, he also observed an enrichment of Phe in GA TMDs and of Ile in PM TMDs.

**Figure 2.**
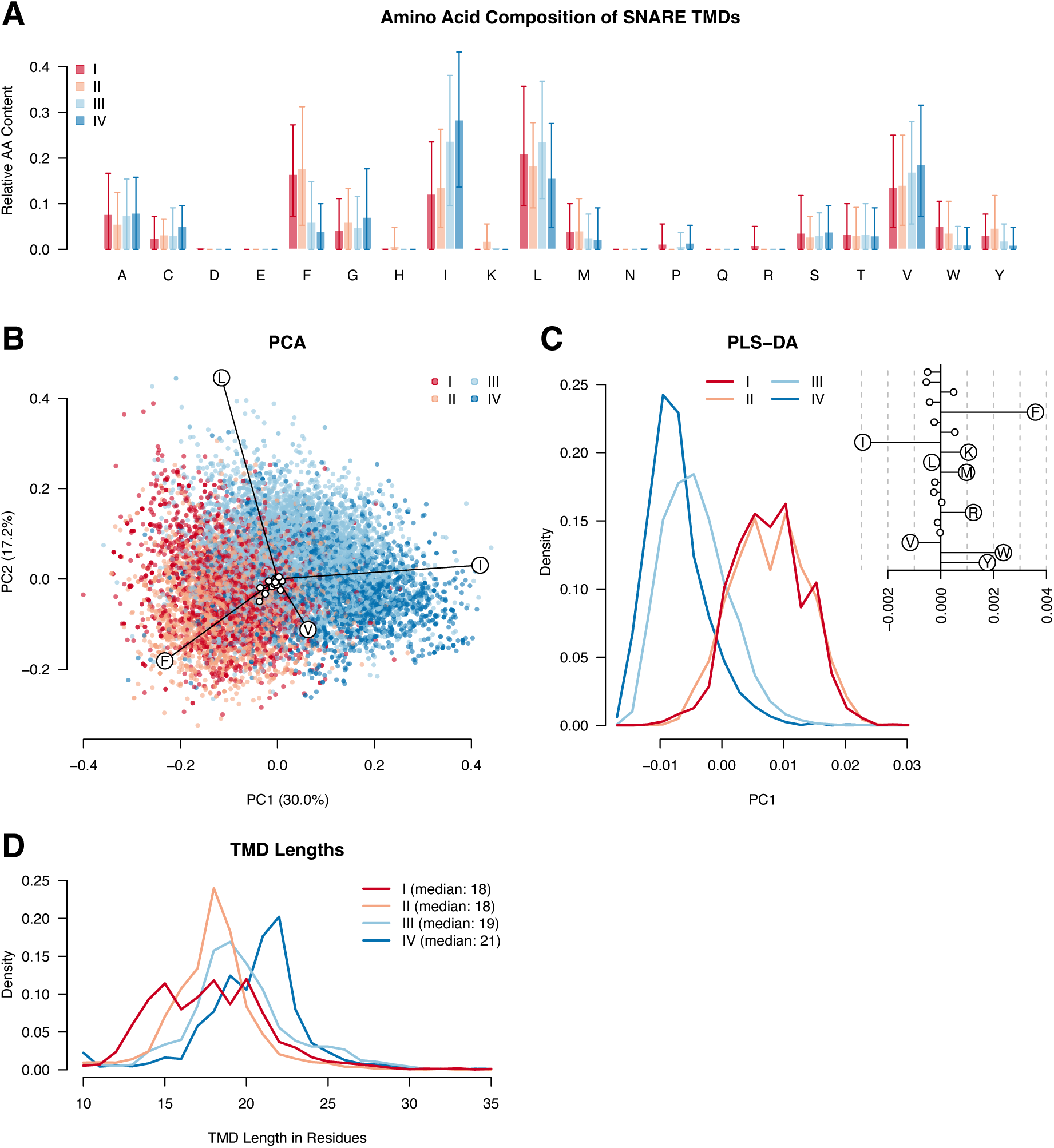
Amino acid composition and length of SNARE TMDs. **(A)** Mean amino acid composition of SNARE TMDs by class. TMDs were identified using the Goldman-Engelman-Steitz (GES) hydrophobicity scale with a rolling mean over 15 residues. Error bars represent the range containing 80% of all TMDs. **(B)** Unscaled principal component analysis (PCA) of SNARE TMD amino acid composition with corresponding relative loadings. Each data point represents a single TMD sequence. A scaled version of this PCA can be found in Fig. S1A and S1B. **(C)** Histogram of partial least squares-discriminant analysis (PLS-DA) principal component 1 (PC1) scores depicting the separation between classes I/II and classes III/IV. Feature weights contributing to the discrimination are shown in the upper-right panel. **(D)** Histogram of TMD lengths of SNAREs calculated using the GES hydrophobicity scale.

Principal component analysis (PCA) reveals a clear compositional dichotomy between classes I/II and III/IV, demonstrating that the split is evident not only in average composition but also at the level of individual TMDs (Fig. 2B). As seen with the averages, the split into the two groups is driven by the differences in Phe and Ile content. Besides these large compositional differences, classes I/II and III/IV also differ in less frequently occurring amino acids. Using a scaled partial least squares-discriminant analysis (PLS-DA) we separated classes I and II from classes III and IV (Fig. 2C). In this analysis, Phe and Ile remain the dominant contributors to the separation, with smaller contributions from additional amino acids being present as well. Aromatic residues (Trp and Tyr) are more abundant in classes I and II, as are Met, Lys, and Arg to a lesser extent. However, positively charged residues are overall rare in TMDs and are only present in a subset of proteins. Specifically, Lys occurs in 33.0% of class II and 5.5% of class I TMDs, whereas Arg is found in 16.9% of class I and 1.5% of class II TMDs. In classes III and IV it is only Val that additionally occurs notably more frequently than in classes I and II. The average Val content continually increases along the secretory pathway.

Although the two dominant compositional groups are evident, the distributions partially overlap. We therefore examined the TMD amino acid composition of all SNAREs individually to see if there were any that could not be easily assigned to either group. Two SNAREs in particular, Gos28 of the Qb II and Nyv1 of the R III subtype, display intermediate properties. The intermediate position stems from a low Phe content in the case of Gos28 and a low Ile content in Nyv1 proteins.

In addition to the clear split between classes I/II and III/IV, the individual SNARE classes can also be separated with PLS-DA, even though a larger overlap remains (Fig. S1C & S1D). Nearly all amino acids contribute to the partial differentiation between classes I and II, including that class II has more Lys and class I more Arg. The separation between classes III and IV is similarly complex, but it is dominated by a higher Leu content in class III TMDs. This higher Leu content in class III is also apparent in the mean amino acid composition (Fig. 2A) and to some degree in the PCA (Fig. 2B). This indicates that although Leu is common in all TMDs and does not contribute to the dichotomy, it helps distinguish classes III and IV.

SNARE TMDs might be influenced by potential variations in membrane compositions between phylogenetic groups. To investigate these potential differences, PCA and PLS-DA were performed separately for metazoan, fungal, archaeplastid, and SAR SNARE TMDs (Fig. S2−S4). Overall, the main compositional dichotomy between classes I/II and III/IV, and the differences among individual classes persist across all phylogenetic groups, but their magnitudes vary. Notably, fungi show a clear separation between classes III and IV, which is evident from PCA alone (Fig. S3E). Similar to the complete dataset, fungi class III TMDs contain more Leu than class IV TMDs, which in turn contains somewhat more Val and Ile. In metazoan SNAREs, the class I vs II split is more distinct than in the other taxonomic groups and shows again the complex contribution from many amino acids (Fig. S3C). The characteristic Ile-Phe dichotomy is conserved across all groups, while the specific amino acids contributing to finer class-level separations differ to some extent among lineages. This suggests that SNARE TMDs may have undergone lineage-dependent adaptations.

In contrast to the dichotomy between early and late secretory pathway, the four basic SNARE types (Qa, Qb, Qc, R) cannot be distinguished by their TMD composition. Both PCA and PLS-DA show that the four SNARE types overlap extensively in their TMD amino acid composition (Fig. S5). In contrast to the TMDs, the composition of the adjacent SNARE motifs is sufficient to distinguish the four SNARE types.^8^ This distinction is expected, as each of the four SNARE types forms a monophyletic group. Differences between the four SNARE types become apparent only when SNAREs are analyzed within a single class (Fig. S6 & S7). These type-specific differences are most pronounced in classes I and II. Taken together, these findings suggest that the amino acid compositions of SNARE TMDs represent independent adaptations within each SNARE type to the distinct biophysical properties of membranes across cellular compartments.

### TMD length

Proteins of the late secretory pathway are thought to have longer TMDs than those of the early secretory pathway, consistent with the increasing thickness of these membranes (Fig. 1D). This finding is largely based on sequence predictions. TMDs are commonly identified from amino acid sequences using hydrophobicity-based scoring methods that estimate whether residues are more likely to reside within the membrane or in aqueous environments.^39^ A wide range of different hydrophobicity scales and online prediction tools are available for this purpose. In their study, Sharpe *et al*. used the Goldman-Engelman-Steitz (GES) hydrophobicity scale to show that TMD lengths of bitopic membrane proteins differ between early and late secretory pathways.^28,40^ Using a GES-based approach, we obtained similar results for our SNARE dataset, with TMD lengths increasing along the secretory pathway (Fig. 2D).

Sequence-based methods delineate hydrophobic stretches but do not necessarily capture the actual membrane-embedded boundaries of TMDs. We therefore asked whether the observed increase in TMD length persists when TMDs are defined by their membrane positioning in all-atom molecular dynamics (MD) simulations. Given the computational cost of all-atom simulations for the complete SNARE repertoire, we limited our analysis to human SNARE TMDs.

Simulations of all human SNARE TMDs were conducted in both DMPC and DOPC membranes, which differ in acyl-chain length and saturation and therefore provide bilayers of distinct thickness (DMPC: ∼36.0 Å, DOPC: ∼38.5 Å considering the phosphate planes) and fluidity. Because membrane-spanning boundaries are not uniquely defined, we quantified TMD lengths using two complementary measures: the number of Cα atoms located between the acyl-oxygen (O–O) or phosphate (P–P) planes of opposing leaflets (Fig. 3A). For each SNARE, TMD lengths fluctuate substantially over time (Fig. 3B & 3C). Nevertheless, replicate simulations and lipid compositions yield largely consistent results, with Pearson correlation coefficients (PCC) between DOPC and DMPC of 0.64 (O–O) and 0.90 (P–P), indicating minimal dependence on the lipid model, particularly for the P–P definition (Fig. S8A & S8B).

**Figure 3.**
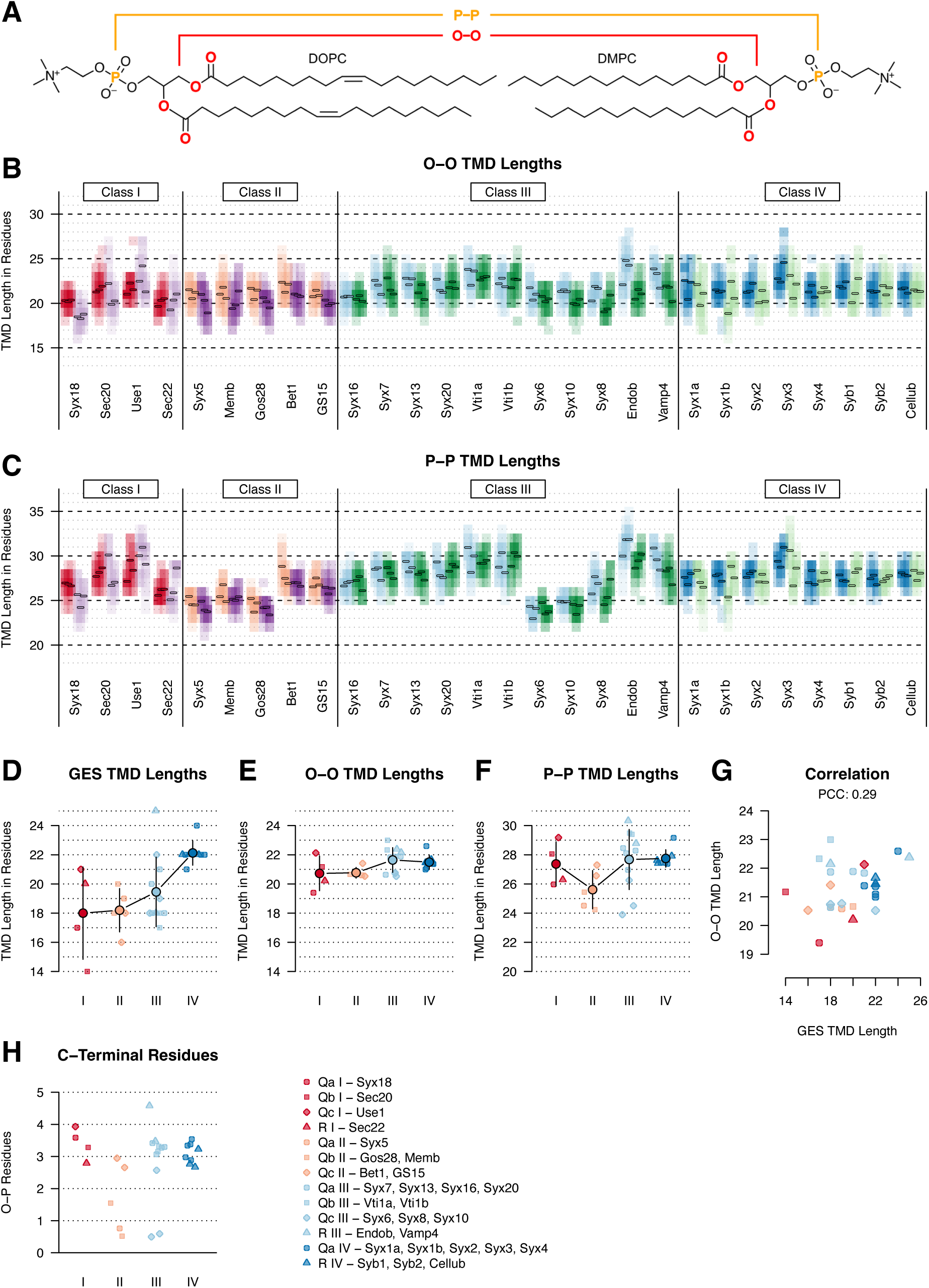
TMD lengths of human SNAREs determined from molecular dynamics (MD) simulations and sequence data. **(A)** Lipid molecules used in the MD simulations. Atoms used to define the membrane boundaries are highlighted in orange (phosphorus atoms for P−P definition) and in red (acyl acid oxygens for O−O definition). **(B)** Histogram of TMD lengths from MD simulations of human SNAREs. Lengths were determined by the Cα atoms located within the O−O layers considering the 10 lipids closest to the TMD in each leaflet. MD simulations were performed in triplicate for each SNARE TMD in DOPC and DMPC bilayers. For each TMD, results obtained with DOPC are shown on the left and those with DMPC on the right using complementary colors. Color intensity indicates how frequently a given TMD length was observed. The average TMD length for each trajectory is shown as a short bar. **(C)** Histogram of human SNARE TMD lengths from MD simulations, based on the number of Cα atoms located within the P−P layers considering the 10 lipids closest to the TMD in each leaflet. Other details are the same as in subfigure B. **(D)** TMD lengths of human SNAREs calculated from sequence data using the GES scale. Values for individual SNAREs are plotted in the background, while class averages are shown in the foreground; bars indicate standard deviations. **(E)** Average TMD lengths from MD simulations using the O–O definition. Individual protein averages are plotted in the background, class averages in the foreground; bars indicate standard deviations. **(F)** Average TMD lengths from MD simulations using the P–P definition. Individual protein averages are plotted in the background, class averages in the foreground; bars indicate standard deviations. **(G)** Correlation between TMD lengths obtained from MD simulations (O–O definition) and those derived from sequence data using the GES hydrophobicity scale. **(H)** Number of residues located between the oxygen and phosphate layers in the outer (luminal or exoplasmic) leaflet. The difference between the O–O and P–P thickness is approximately 8.5 Å, corresponding to about 5.5 residues in an α-helix (1.5 residues per 1 Å). Roughly half of this distance (∼3 residues) is expected to lie between the O and P layers. Smaller numbers indicate that the *C*-terminus does not reach the phosphate layer.

Using the O–O criterion, TMD lengths fall mostly between 20 and 25 residues, with little variation among SNAREs (Fig. 3B). Greater heterogeneity is observed with the P–P definition, where most SNARE TMDs span 25 to 30 residues, but five proteins (Syx5, Memb, Gos28, Syx6, and Syx10) exhibit noticeably shorter apparent lengths (Fig. 3C). This reduction arises because their *C*-termini fail to fully traverse the distance between the oxygen and phosphate planes at the outer (luminal) leaflet, resulting in fewer residues between these planes compared with other SNAREs (Fig. 3H, S8C & S8D).

In contrast to the sequence-based GES analysis (Fig. 3D), the trend of increasing TMD length along the secretory pathway is markedly attenuated in the MD simulations. Average O–O-based lengths increase only marginally across classes I–IV (Fig. 3E), remaining well within the intrinsic fluctuations observed during the simulations and substantially smaller than the differences inferred from GES. A similar pattern is observed for the P–P definition, where the shorter average length of class II TMDs is driven by three proteins with short *C*-termini (Fig. 3F). Excluding these proteins aligns class II more closely with class I, while class III TMDs become slightly longer than those of class IV (27.4, 26.9, 28.4, and 27.7 residues for classes I–IV). A graphical overview of the various TMD segments identified with each method for human SNAREs is provided in figure S9.

The discrepancy between GES- and MD-derived TMD lengths is further evident in the weak correlation between them: The PCC is 0.29 between GES and O–O TMD lengths (Fig. 3G) and 0.28 between GES and P–P TMD lengths (Fig. S8E). This creates a potential issue for the statistics on TMD amino acid composition we presented in the previous section, where we aimed to analyze the composition of the membrane-embedded portion of the protein. If TMD boundaries are not accurately captured with GES, then the amino acid statistics could be considerably different with the MD-defined boundaries. To evaluate this effect, amino acid compositions of human SNARE TMDs were recalculated. Amino acid contents remain similar overall between GES and O–O TMDs, whereas larger differences occur with P–P TMDs (Fig. S10A). This is consistent with the smaller length differences between GES and O–O TMDs and the considerably longer spans of the P−P TMDs. The longer P–P spans include regions flanking the hydrophobic core, leading to a general decrease in hydrophobic amino acid content and an increase in positively charged residues.

Similarly, PLS-DA scores were recalculated for the human SNARE TMDs to assess the impact of the different TMD definitions (Fig. S10B–S10D). Although the separation between classes I/II and III/IV becomes less pronounced, especially for P–P TMDs, the scores remain strongly correlated across methods (PCC = 0.82–0.89). Interestingly, Gos28 becomes even more similar to class III/IV TMDs with the O–O criterion than was the case with GES. However, because the PLS-DA model was trained on GES-defined TMDs, recalibrating it with MD-based TMDs could improve the distinction between classes I/II and III/IV again.

### Lipid-accessible surface area

TMDs of ER and GA SNAREs differ from those in the TGN, ES, SV, and PM in their Phe and Ile content. Both amino acids are among the most hydrophobic residues, but they differ in side-chain architecture: Phe carries a bulkier side chain than Ile, providing a larger surface for interactions with surrounding lipids (Fig. 4A). However, side-chain geometry alone can play an important role: Despite differing only by the position of a single methyl group, Leu generates a larger interaction surface than Ile because the γ-branch in Leu positions its methyl group farther from the protein backbone (Fig. 4A).^41^

**Figure 4.**
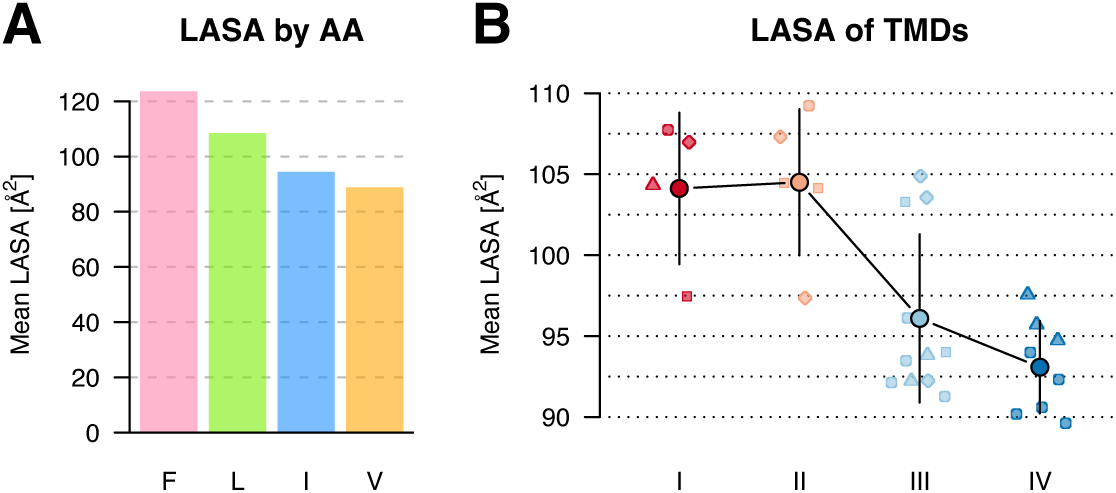
Lipid-accessible surface area (LASA) of human SNAREs. **(A)** Mean LASA of Phe, Ile, Leu, and Val within a model TMD. Twenty residues in the TMD of Syb2 (M29–F48, Alignment S1) were mutated to each of the four amino acids. The bar plot shows the average LASA per residue within the mutated span. **(B)** Average LASA per residue of human SNARE TMDs. Average values of individual SNAREs are shown in the background, class averages are displayed in the foreground, and error bars indicate the standard deviation across individual SNAREs.

To determine whether the compositional differences translate into altered lipid interaction surfaces, we calculated the LASA for human SNARE TMDs (Fig. 4B). Indeed, the average LASA per residue calculated from MD simulations using the O–O definition shows a pronounced difference between classes I/II and III/IV, with classes I and II exhibiting the larger lipid-interacting surfaces. Some exceptions exist: Sec20 (class I) and GS15 (class II) show markedly smaller LASAs, whereas Vti1b, Syx6, and Syx10 (all class III) display larger values, all of which resemble more closely the values of the opposite group. Notably, Syx6 and Syx10 are also among the TMDs with shorter lengths.

In the human SNARE TMD set, and considering the four major hydrophobic amino acids, Phe and Ile contents correlate most strongly with LASA, indicating that these residues are the primary drivers to control lipid accessibility (Fig. S11A–S11D). Accordingly, the PLS-DA scores for amino acid composition closely mirror the LASA values (Fig. S11E). Some of the largest deviations occur in the five TMDs with unexpectedly large or small LASA and, notably, in Gos28. Although its PLS-DA score places Gos28 closer to class III/IV SNARE TMDs in terms of composition, its LASA remains large and comparable to that of other class I/II members. This elevated LASA is driven at least in part by its high Leu content, a pattern also observed in Vti1b, Syx6, and Syx10, all of which contain relatively few Phe residues and appear to achieve a large LASA through increased Leu levels.

### C-terminal ends of TMDs

In most human SNAREs, the TMD extends no more than a single residue beyond the luminal phosphate plane of the membrane (Fig. S8D). In some cases, the TMD is shorter and ends before reaching the phosphate plane, as mentioned above, whereas in a few proteins it extends beyond this boundary, such as in Sec20, Gs15, Use1 and Vti1a. Among the latter, Sec20 exhibits the largest number of residues extending beyond the membrane with an average of four. The Sec20 *C*-terminus found in the human protein is highly conserved within animals of the deuterostome lineage but varies across eukaryotic phyla. In many plants and fungi, the Sec20 *C*-terminus is substantially longer, extending beyond 100 residues, and contains a KDEL signal or related motif such as RDEL or HDEL for ER retention.^14^

The two longest *C*-termini in human SNAREs are found in Vamp7 and Myob, which were not included in the analysis because their extended *C*-termini partially insert into the membrane during the MD simulations, thereby interfering with the measurements. The *C*-terminus of human Myob contains roughly 22 residues and is most likely intrinsically disordered based on its sequence. Similar luminal *C*-terminal extensions are also found in several non-human R IV proteins. Vamp7 possesses a highly conserved *C*-terminal region that AlphaFold 2 (AF2) predicts will fold back on itself and form a disulfide bond (Fig. 5H). This curled *C*-terminus is not only present in the human ortholog but is highly conserved across all major eukaryotic lineages, suggesting that it represents the ancestral state already present in the LECA (Fig. S12).

**Figure 5.**
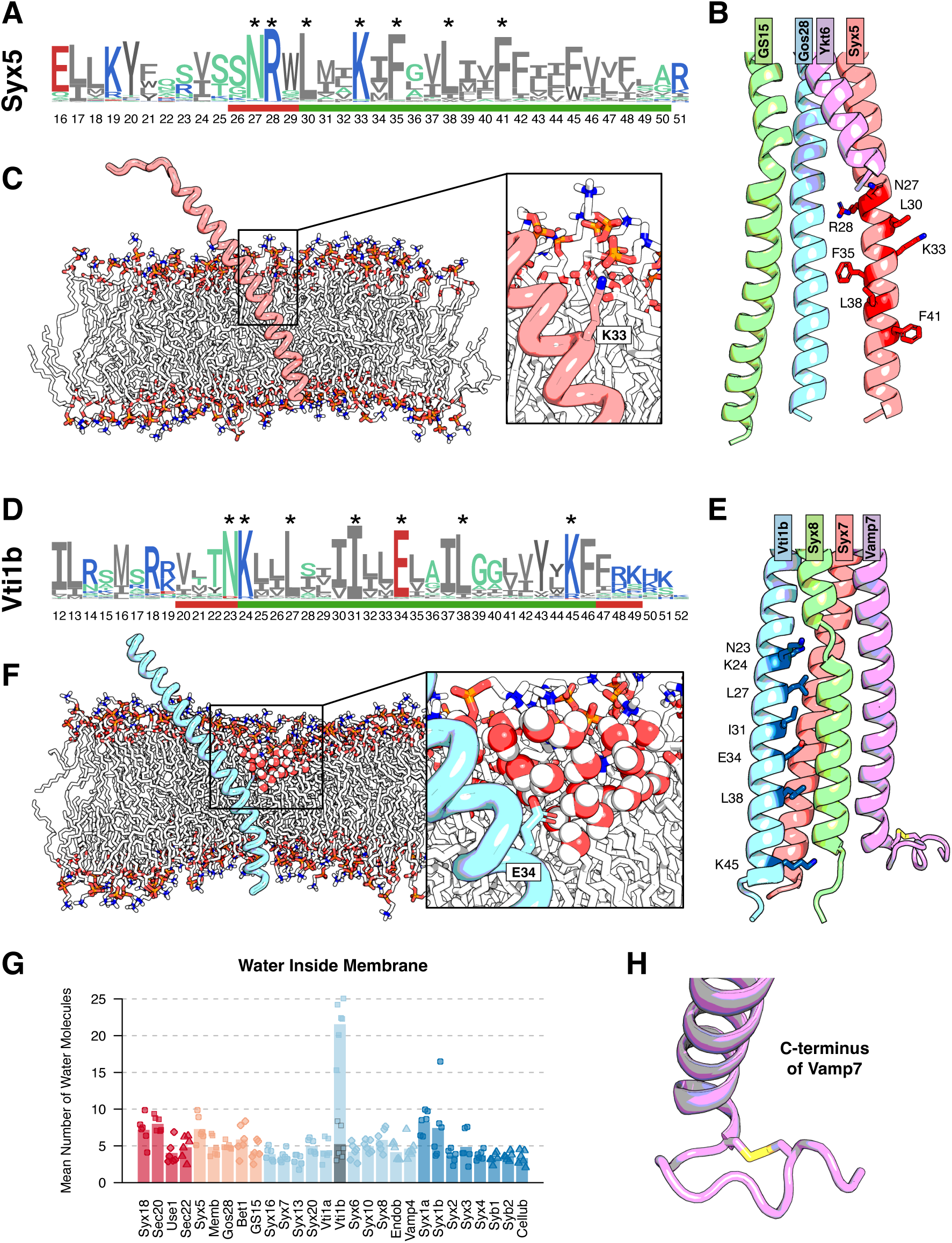
Potentially charged residues within Syx5 and Vti1b TMDs. **(A)** Sequence logo of Syx5 based on 652 sequences. Stars mark the seven most conserved residues in the TMD. Red and green bars indicate the TMD span based on the P–P and O–O definitions, respectively. The *C*-terminal residue is not present in human Syx5. Amino acids are color-coded according to their chemical properties. Sequence numbering corresponds to the models used in MD simulations, which were based on the last 50 residues of the human SNAREs. **(B)** AF2 model of the tetramer formed by human Syx5, Gos28, GS15, and Ykt6. Side chains of the seven most conserved TMD residues in Syx5 are highlighted. Ykt6 lacks a TMD and is membrane-anchored via palmitoylation. **(C)** MD snapshot of Syx5 embedded in a DOPC bilayer. The charged residue K33 is positioned within the membrane, but its side chain extends towards lipid phosphates and headgroups. **(D)** Sequence logo of Vti1b based on 154 sequences. Stars mark the seven most conserved residues in the TMD. Red and green bars indicate the TMD span based on the P–P and O–O definitions, respectively. Amino acids are color-coded according to their chemical properties. Sequence numbering corresponds to the models used in MD simulations, which were based on the last 50 residues of the human SNAREs. **(E)** AF2 model of the tetramer formed by human Syx7, Vti1b, Syx8, and Vamp7. The seven most conserved residues in Vti1b are highlighted. The AF2 model predicts a potential disulfide bond at the *C*-terminus of Vamp7. **(F)** MD snapshot of Vti1b in a DOPC bilayer. The buried charged residue E34 induces the formation of a water pocket within the membrane. **(G)** Number of water molecules within the membrane based on the O–O definition. Only water molecules within 10 Å (in the x-y plane) around the TMD center were considered. Bars represent protein averages; data points show averages from individual trajectories. The grey Vti1b bar indicates the result for the protonated (and therefore neutral) E34. **(H)** AF2 model of the *C*-terminal end of human Vamp7. The predicted disulfide bond is highlighted in yellow.

### Conserved charges in TMDs

Our sequence analysis shows that potentially charged residues (Arg, Lys, His, Asp, Glu) are rare within SNARE TMDs but are slightly enriched in ER and GA SNARE TMDs, where they are present at very low levels (Fig. 2A). Using the GES-based boundaries, only four of the 28 investigated human SNAREs contain a single potentially charged residue within their TMD: Sec22 (Arg), Syx5 (Lys), Gos28 (His), and Vti1b (Glu). While the His in Gos28 is located at the *C*-terminus, the charged residues in the other SNAREs lie within the hydrophobic core of the TMDs, with the Glu in Vti1b positioned at its very center. Whereas the potentially charged residues are highly conserved in Syx5 and Vti1b, the situation is a bit more complex in the other two proteins. The *C*-terminus of Gos28 is most often formed by a potentially charged residue, which is typically Arg or Lys outside the animal lineage of Deuterostomia, within which it is predominantly His. Similarly, most Sec22 proteins contain either a Lys or Arg residue towards the *C*-terminal end of the TMD. However, the exact position of this potentially charged residue does not appear to be conserved.

The membrane interior is highly hydrophobic, disfavoring polar or charged groups unless they can form compensating stabilizing interactions. Thus, the presence of even a few potentially charged residues within SNARE TMDs is surprising and raises the question of how they are accommodated within the hydrophobic environment of the membrane. Using snapshots from the MD simulations of the monomers and AF2 predictions of the tetramers, we took a closer look at the potentially charged residues in Syx5 and Vti1b.

Syx5 contains a Lys within its TMD that is highly conserved across Eukaryota (Fig. 5A). In addition to this Lys, several other residues are strongly conserved, but they do not align on the same face of the helix as can be seen in a model of the Golgi SNARE complex, indicating that they are unlikely to form a shared interaction surface (Fig. 5B). The long aliphatic chain of Lys allows its Cα to remain buried in the membrane while the terminal amino group reaches and interacts with the polar headgroup region of the lipids, a conformation commonly referred to as “snorkeling” (Fig. 5C).^42^ It is therefore plausible that this Lys remains positively charged in the membrane and that it has a role in restricting the orientations that the Syx5 TMD can adopt. This interpretation is further supported by the observation that substitution of Lys with Leu promotes deeper insertion of the TMD into the membrane, resulting in partial membrane insertion of several residues flanking the TMD at the *N*-terminal side (Fig. S13). Interestingly, few exceptions exist in which Lys is substituted by Gln, such as in *Plasmodium* and *Caenorhabditis* (including *C. elegans*) Syx5, which might fulfill a similar functional role.

The transition to multicellularity at the base of Metazoa was accompanied by an expansion of the set of endosomal SNAREs through the emergence of new paralogs.^2^ Vti1b is such a metazoan SNARE and contains a highly conserved Glu within its TMD (Fig. 5D). Unlike Syx5, the seven most conserved residues in Vti1b align on the same face of the helix, likely forming the TMD-TMD interaction surface in the tetrameric complex (Fig. 5E). Consequently, the potentially charged Glu residue is buried within the tetrameric core, where it may interact with backbone amides, as no complementary charged or polar side chains are expected nearby.

While the tetrameric arrangement poses no issue for a charged Glu, the situation is different in the monomeric Vti1b (Fig. 5F). In the absence of a stabilizing partner, the charged Glu side chain induces the formation of a water pocket within the membrane interior. No other SNARE TMD produced similar water-filled cavities in the MD simulations (Fig. 5G). However, the low dielectric constant of the membrane interior can substantially shift the pKa values of ionizable residues, potentially allowing their side-chain charges to be neutralized even at physiological pH.^43^ When Glu is protonated, and thus neutral, no water pocket forms in MD simulations. The protonation does not affect TMD length (Fig. S14), suggesting that other residues, particularly positively charged amino acids near both TMD flanks, determine the effective span. The strong membrane deformation by charged Glu suggests that the Glu residue may be protonated and thus neutral *in vivo* or that an interaction partner provides a compensating charge in the native membrane.

TMD boundaries are commonly assigned based on the position of charged residues in sequence-based prediction, even though such residues can be partially or fully buried within the membrane, as illustrated for Syx5 and Vti1b. Based on sequence alone, only 4 of the 28 analyzed human SNAREs contain potentially charged residues within their TMD. This assessment changes markedly when MD-derived boundaries are considered. Using the O–O definition, the TMD is frequently shifted toward the *N*-terminus relative to the GES-defined TMD, resulting in the inclusion of additional *N*-terminal residues and the exclusion of *C*-terminal residues (Fig. S9). As a result, 80% of the analyzed human SNAREs contain at least one potentially charged residue within the membrane-embedded segment using the O–O criterion. Syx5 is therefore not an outlier with its Lys but instead displays a common feature among SNARE TMDs. The potentially charged residues present in O–O TMDs occur close to the cytosolic boundary and are positioned one to three residues closer to the membrane-water interface than in Syx5. Even more charged residues are included with the P–P definition because this boundary encompasses additional flanking residues. Under this definition, all TMDs contain at least two potentially charged residues.

## Discussion

The starting point of this study was the question of whether physicochemical differences between intracellular membranes are reflected in the amino acid composition of SNARE TMDs. Our analysis reveals that SNARE TMDs segregate into two broad compositional groups that correlate with their subcellular localization. SNAREs functioning at the ER and Golgi apparatus are enriched in Phe, whereas those acting at the *trans*-Golgi network, the endosomal compartments, secretory vesicles and the plasma membrane contain higher levels of Ile, with smaller contributions from additional residues. Importantly, this dichotomy is evident not only in average compositions but also at the level of individual TMDs. A difference in Phe and Ile composition between GA and PM TMDs was reported previously by Sean Munro based on a limited dataset of various proteins.^38^ Our study extends this principle to a larger TMD collection and shows that such compositional biases are a robust feature within the SNARE family across the eukaryotic endomembrane system. In addition to the work by Sean Munro, several other studies have reported related observations on TMD amino acid composition and cellular localization, including analyses that specifically addressed SNAREs.^17,33,44–46^ Collectively, these observations are consistent with the patterns identified in our dataset.

We next asked how these compositional differences relate to subcellular localization: do they represent adaptations to the physicochemical properties of the native membranes, or do they actively contribute to protein sorting? Proper SNARE localization is essential, as only correctly distributed SNAREs can assemble into productive fusion complexes. Because SNAREs reside in the fused membrane after they catalyze membrane fusion, they must be actively trafficked back to their original compartments to enable subsequent fusion events. Thus, SNARE distribution must be continuously maintained by retrieval and re-sorting mechanisms, making their localization not static but the result of ongoing sorting.

SNARE distribution is governed by multiple determinants. These include protein-protein interactions with SM proteins, tethering factors, sorting receptors and coat complexes through defined sorting signals in either soluble domains or the TMD.^12,13^ For example, the *C*-terminal KDEL motif of Sec20 mediates retrieval to the ER, and a dileucine-based motif in Vamp4 likely contributes to its trafficking itinerary.^14,47^

Beyond these canonical sorting determinants, the physicochemical properties of the TMD itself may contribute to localization.^17,48^ The compositional dichotomy we observe in SNARE TMDs is consistent with such a role. However, these differences could also simply reflect energetically favorable protein-lipid interactions within distinct membrane environments.^41,44^ Along the secretory pathway, membrane rigidity increases, with ER membranes being the most loosely packed and deformable, whereas the PM is the most rigid and tightly packed (Fig. 1E). Loosely packed membranes contain more packing defects, favoring extensive protein-lipid interactions. In contrast, tight lipid packing in membranes such as the PM disfavors such interactions, as they would disrupt lipid-lipid contacts. Consequently, TMDs in rigid membranes may minimize their hydrophobic lipid contact area through a reduction in LASA, whereas TMDs in more deformable membranes may benefit from maximizing it.

Importantly, these membrane adaptation and sorting through TMD properties are potentially linked phenomena.^38,44^ At the TGN, lipid microdomains (also referred to as lipid rafts) have been proposed to contribute to the sorting of a subset of PM-directed proteins.^49–52^ TMDs with low LASAs preferentially partition into these lipid microdomains, whereas specific residues such as Phe and Leu can reduce raft affinity and thereby influence trafficking routes.^46,53^ Based on the calculated LASA values for human SNAREs, raft affinity appears to be a common feature of class III (TGN, ES) and class IV (SVs, PM) SNAREs, suggesting that TMD-dependent partitioning could support their forward trafficking. Although this tendency is not evident from average LASA values alone, the higher Leu content in class III SNAREs may partially counteract raft partitioning and thereby restrict their steady-state localization to endosomal compartments.

Notably, several human SNARE TMDs deviate from the general LASA-based trends, arguing against a model in which TMD properties alone dictate localization. Indeed, multiple sorting determinants can coexist within a single protein. For example, the yeast SNAREs Sed5 and Ufe1 combine TMD features with cytosolic signals, likely enabling more precise trafficking control.^45,54^ Against this background, the compositional differences observed in SNARE TMDs are best interpreted as contributing to, but generally not being sufficient for, compartment-specific localization.

Another parameter frequently discussed in the context of membrane protein sorting is TMD length, which is thought to increase by four to nine residues along the secretory pathway, in line with the increase in membrane thickness.^16,28,33,38,44^ According to the hydrophobic matching hypothesis, TMDs preferentially partition into membranes matching their hydrophobic length.^18,28^ This sorting mechanism was suggested to act in lipid rafts, which have thicker membranes than the surrounding disordered region and are thought to enrich long TMDs for PM trafficking.^50,53^

Using sequence-based hydrophobicity predictions, we observe a gradual increase in predicted TMD length along the secretory pathway, both in the expanded dataset and when considering human SNAREs alone. However, when TMD boundaries are defined by membrane positioning in all-atom MD simulations, this gradient becomes far less pronounced. This discrepancy reflects fundamental differences between the two approaches.

Sequence-based methods primarily delineate the most hydrophobic stretch of a transmembrane segment. Charged residues are commonly used to define TMD boundaries and are therefore typically assigned to extra-membranous regions. However, such assignments do not necessarily reflect their actual positioning within the bilayer. For example, Lys is strongly hydrophilic according to hydropathy scales, yet it contains a long aliphatic side chain that can be accommodated into the hydrophobic region of the membrane while its charged ammonium group remains in contact with lipid phosphates or water. Moreover, lipid bilayers do not form sharp boundaries between aqueous and hydrophobic regions but instead contain interfacial transition zones enriched in polar and charged lipid moieties. Additionally, both proteins and membranes are dynamic, and fluctuations further blur static boundary definitions.

Consistent with this view, MD-derived TMD boundaries frequently differ from the hydrophobic stretch determined by sequence analysis. The more stringent O–O definition emphasizes the hydrophobic core, whereas the P–P definition includes more of the interfacial region and therefore incorporates additional polar or charged residues into the membrane-embedded segment. The O–O definition yields highly overlapping TMD length distributions across SNAREs, indicating that the membrane-embedded segments match the same range of membrane thicknesses. With the P–P definition, most short TMDs are found in the GA, which was also observed for fungal TMDs by Sharpe *et al*. and aligns with experimental results that indicate that GA membranes are thinner than ER membranes.^28,55^ Importantly, under both definitions, differences in effective membrane-spanning length between SNARE classes are modest at best compared with the compositional differences described above. These observations indicate that sequence-based TMD-length determination might primarily capture differences in composition rather than substantial differences in membrane-embedded spans.

The MD simulations also help distinguish between two conceptually different types of charged residues within SNARE TMDs. In Syx5, a Lys residue is positioned close to the membrane-water interface. In the simulations, its aliphatic portion remains embedded while the charged group snorkels toward the polar headgroup region. This positioning appears to contribute to a defined alignment of the helix within the membrane, suggesting a structural role in orienting the TMD.

In contrast, Vti1b contains a Glu residue located centrally within the hydrophobic core of the TMD. Unlike interfacial Lys, this residue cannot reach the membrane surface and instead induces local rearrangements of surrounding lipids and water penetration in the simulations when charged. Comparable centrally buried charged residues have been described in other fusion-related membrane proteins, including the HIV-1 envelope protein and the mitochondrial fusion protein Fzo1, where they modulate helix-helix interactions, lipid packing, and water retention inside the membrane.^56,57^ In case of Fzo1, the packing defects expose hydrophobic lipid tails to the solvent, which is thought to facilitate membrane fusion. By analogy, the conserved Glu in Vti1b may influence TMD organization, SNARE complex assembly, or local membrane properties during fusion. The restricted occurrence of such central charges among SNAREs suggests specialized roles rather than a general principle. At the same time, charged residues within TMDs have been implicated in ER retention and degradation pathways, indicating that such features may also affect localization and turnover.^58^

Taken together, our results show that SNARE TMDs exhibit systematic differences in amino acid composition that correlate with their site of action, whereas differences in effective membrane-spanning length are small or non-existing when membrane embedding is assessed dynamically. By focusing on a single protein family with a conserved core function, our analysis minimizes confounding effects from unrelated evolutionary pressures and highlights a common principle: SNARE TMDs appear tuned to their membrane environments primarily through compositional biases that influence protein-lipid interactions, while length variations play a comparatively minor role. We therefore propose that TMD composition contributes to compartmental localization and functional optimization by modulating energetic compatibility with specific bilayer environments, in concert with protein-based sorting mechanisms.

## Methods

All SNARE sequences were obtained from the TRACEY database,^1,37^ yielding a total of 18,810 sequences. TMDs were identified using the GES hydrophobicity scale, which specifically accounts for α-helical structures.^40^ Hydropathy scores were calculated using a weighted rolling average with a 15-residue window: the central residue was fully weighted, while flanking residues were linearly downweighted, reaching zero weight seven positions away from the center. Only TMDs located *C*-terminally to the SNARE domain and between 10 and 35 residues in length were retained for further analysis. The TMDs of Syx17, which contain two α-helices, were excluded. This filtering step yielded a total of 13,941 TMD sequences. Amino acid compositions were analyzed using PCA with centering but without scaling, and PLS-DA with both centering and scaling. PLS-DA is a supervised dimensionality reduction method that incorporates class labels to enhance group separation and is often considered a supervised analogue of PCA.^59^

Because SNARE proteins are represented in unequal numbers in the dataset, potential sampling bias was evaluated. To test whether the observed compositional differences reflect true class-dependent variation, we performed a more stringent analysis (Fig. S15). In this analysis, only classes I and III were compared, as these are the only classes containing TMDs from all four SNARE types (Qa, Qb, Qc, and R). In addition, only unique TMD sequences were included. To eliminate sample-size bias, 1,000 TMDs were randomly drawn (with replacement) from each type-class combination (for example, Qa II SNAREs). Results from this reduced dataset closely match those of the full analysis, indicating that the findings are robust. The largest differences in PLS-DA weights occur for less frequent amino acids, notably Lys and Tyr, which are characteristic of class II but rare in class I. Nonetheless, the distinction between classes I and III remains clearly defined in both PCA (Fig. S15C) and PLS-DA (Fig. S15D), with Phe and Ile again serving as the main discriminants.

MD simulations were based on structural models for SNAREs made by ColabFold: AlphaFold2 using MMseqs2.^60^ Only the *C*-terminal 50 residues of human SNAREs were used to create the structural models (Alignment S1). The only exception was Myob, which was truncated after the TMD due to its extended *C*-terminus, resulting in a construct of 55 residues. AF2 models for Syx8 and Syx10 displayed pronounced kinks between the SNARE domain and the TMD. These distortions were manually corrected in PyMOL 3.0.5^61^ to prevent the *N*-terminus from entering the membrane.

The thirty human SNAREs containing a single TMD were embedded in lipid bilayers using Membrane Builder^62,63^ in CHARMM-GUI.^64–66^ Protein charges were kept at their default protonation states in CHARMM-GUI, corresponding to standard ionization at neutral pH. For Vti1b, a separate simulation was performed in which the Glu residue within the TMD was protonated. Each system was solvated with 25 Å minimum water height. Bilayers consisted of either DOPC or DMPC lipids (100 lipids per leaflet) arranged in a hexagonal box. Protein orientation was determined using PPM2,^67^ which successfully positioned most proteins except Vti1b, Syx6, and Endob. For these three proteins, orientation was manually defined by aligning a vector along the z-axis between the following residue pairs: Vti1b (L26–F46), Syx6 (W30–V49), and Endob (M26–F44).

For every protein-lipid setup, three independent simulations were conducted, consisting of CHARMM-GUI default energy minimization and equilibration steps, followed by a 100 ns production run using GROMACS (2024.1-spack)^68^ at 310 K and CHARMM-GUI default parameters. Only the second half (51–100 ns) of each trajectory was used for analysis.

MD TMD lengths were calculated based on the 10 lipids closest to the helix in each leaflet. Lipid adjacency was determined by the Cα-phosphate distance in three dimensions. The phosphorus atoms and acyl oxygens of the selected lipids were used as boundaries to define whether a residue (based on its Cα position) was embedded within the membrane, considering the z-dimension only. These boundaries are referred to as P–P and O–O. To minimize noise from local fluctuations, a rolling average over five residues was applied to smooth the Cα-phosphorus and Cα-oxygen distance profiles. TMD lengths were calculated as the span from the first to the last residue within the boundaries; therefore, they are not always identical to a simple count of Cα atoms within the boundaries in rare cases when a subsequent Cα atom lies outside the boundary again.

The TMDs of Vamp7 and Myob were excluded from the analysis because both proteins contain an extended *C*-terminal tail. In the case of Vamp7, the tail may adopt a defined fold stabilized by a disulfide bond, as suggested by AF2 (Fig. 5H). This tail resides at the membrane-solvent interface and occasionally inserts into the bilayer. Myob contains a long, relatively hydrophobic tail that begins to penetrate the membrane during MD simulations. In both proteins, TMD lengths fluctuate substantially, making them unsuitable for analysis.

LASAs were calculated from the last frame of each trajectory using a probe radius of 1.88 Å, corresponding to the size of a methylene group within a lipid acyl chain.^41^ LASAs were computed for TMDs as defined by the O–O criterion. Residues located within the membrane in at least 50% of all frames were classified as TMD residues. In the case of the Syb2 mutants, LASAs were calculated based on the last frame of MD simulations with a 1 ns production run in a bilayer containing 50 DOPC molecules in each leaflet.

Water molecules within the membrane were quantified using the O–O boundaries, considering a 10 Å radius in the x-y plane around the Cα atom at the membrane center. All computational analyses were performed in R 4.4.1^69^ using RStudio^70^ with the following packages: bio3d^71^, ggplot2^72^, ggseqlogo^73^, mdatools^74^, msa^75^, and vanddraabe^76^. Protein structures were visualized in PyMOL 3.0.5^61^, sequences were aligned using AliView 1.28^77^ and molecular structures were drawn using ChemDoodle (Version 12.9.0).

## Acknowledgements

This work was supported by the Swiss National Science Foundation (SNF 182732 and SNF 219549). We thank Wan-Chin Chiang and Sévan Stroh for input on the manuscript. ChatGPT and Microsoft Copilot were used to improve the language and ChatGPT was used to redraw the outlines of the hand-drawn sketch of the cell schematic shown in the graphical abstract and Figure 1B.

## Data availability statement

The data that support the findings of this study will be made openly available in the Tracey database in an upcoming publication. This database is a collection of sequences from different protein families, including SNARE, AAA and Rab, classified by HMM profiles.^1,37^

## CRediT authorship contribution statement

**Christian Baumann:** Conceptualization, Methodology, Formal analysis, Investigation, Writing - Original Draft, Visualization **Carlos Pulido Quetglas:** Resources, Data Curation **Dirk Fasshauer:** Writing - Original Draft, Supervision, Funding acquisition.

## Abbreviations

AF2: AlphaFold 2
DMPC: 1,2-dimyristoyl-sn-glycero-3-phosphocholine
DOPC: 1,2-dioleoyl-sn-glycero-3-phosphocholine
ER: endoplasmic reticulum
ES: endosomal system
GA: Golgi apparatus
GES hydrophobicity scale: Goldman-Engelman-Steitz hydrophobicity scale
GET pathway: guided entry of tail-anchored proteins pathway
LASA: lipid-accessible surface area
LECA: last eukaryotic common ancestor
MD: molecular dynamics
O–O: oxygen-to-oxygen membrane thickness definition
P–P: phosphate-to-phosphate membrane thickness definition
PCA: principal component analysis
PC1: principal component 1
PCC: Pearson correlation coefficient
PLS-DA: partial least squares discriminant analysis
PM: plasma membrane
SM proteins: Sec1/Munc18 proteins
SNARE: soluble *N*-ethylmaleimide-sensitive factor attachment protein receptor
SV: secretory vesicle
TGN: *trans*-Golgi network
TMD: transmembrane domain

