## supplementary information for "Transmembrane domain composition reflects subcellular localization of SNARE proteins"

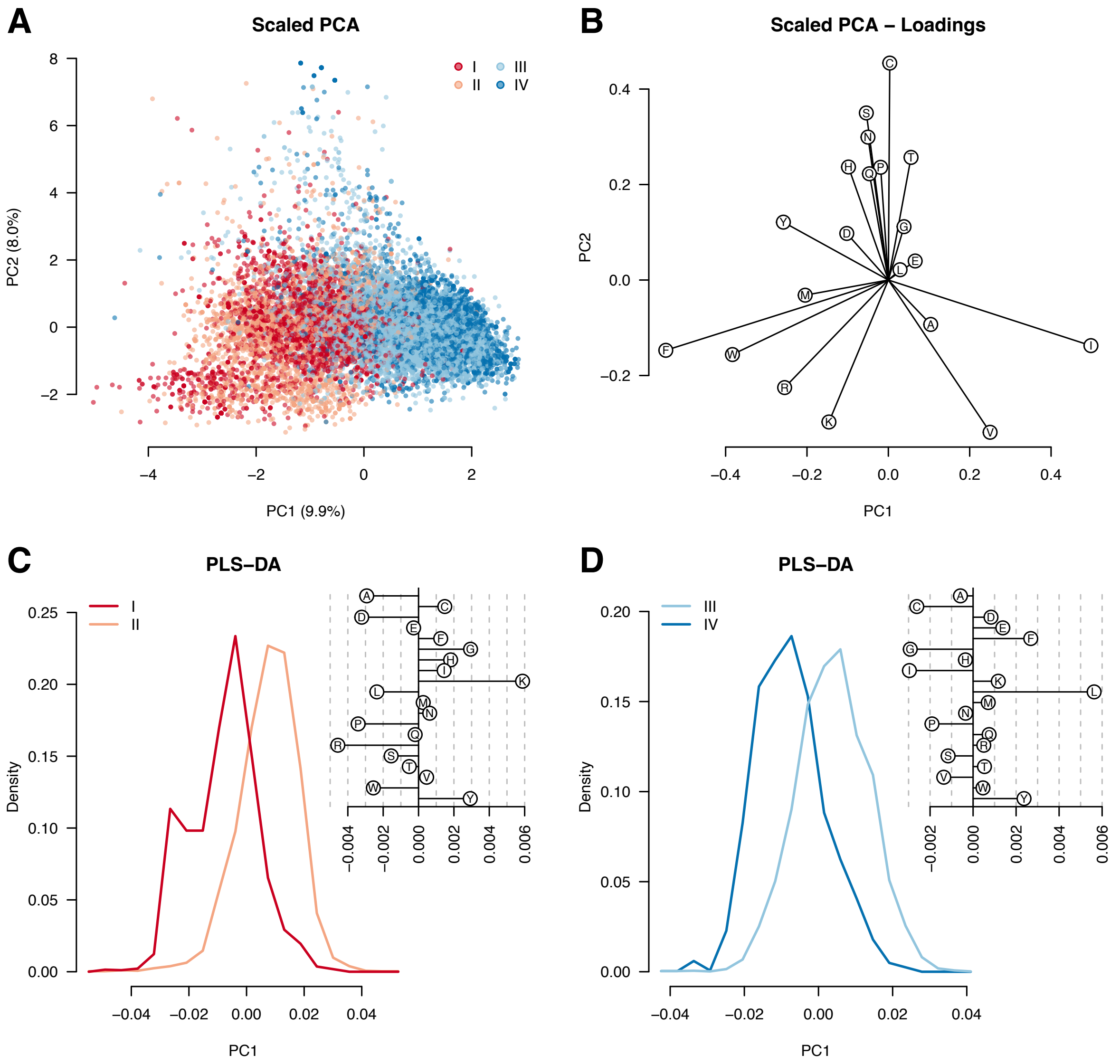


**Figure S1.** PCA and PLS-DA of SNARE TMD amino acid composition. **(A)** PCA of SNARE TMD amino acid composition (centered and scaled). Each point represents one TMD sequence. Class I scores below −1 in both dimensions stem mostly (87.7%) from the R-SNARE Sec22. **(B)** Corresponding PCA loadings. **(C)** Histogram of scores from PLS-DA separating classes I and II. The associated feature weights are shown in the upper right. The minor class I peak is formed by an excess (72.0%) of Sec22. **(D)** Histogram of scores from PLS-DA separating classes III and IV. The associated feature weights are shown in the upper right.

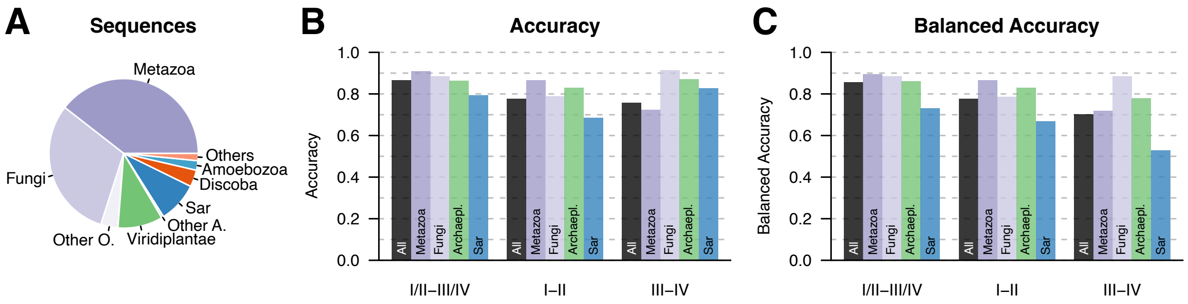


**Figure S2.** Phylogenetic distribution and classification accuracy of SNARE TMDs. **(A)** Number of SNARE TMD sequences per phylogenetic group. Opisthokonta are shown in purple and Archaeplastida in green. SNARE TMDs not belonging to the major groups within Opisthokonta and Archaeplastida are binned in "Other O." and "Other A.", respectively. The number of sequences is: Metazoa (5503), Fungi (4263), other Opisthokonta (519), Viridiplantae (1358), Rhodophyta (46), Glaucocystophyceae (18), SAR (1222), Discoba (507), Amoebozoa (286), Metamonada (120), Haptista (41), Apusozoa (28), Cryptophyceae (21), Malawimonadida (9). **(B)** Classification accuracy of the PLS-DA models shown in Figures 2, S3, and S4. **(C)** Balanced accuracy of the same models. Balanced accuracy was calculated as the mean of sensitivity and specificity, to account for unequal class sizes.

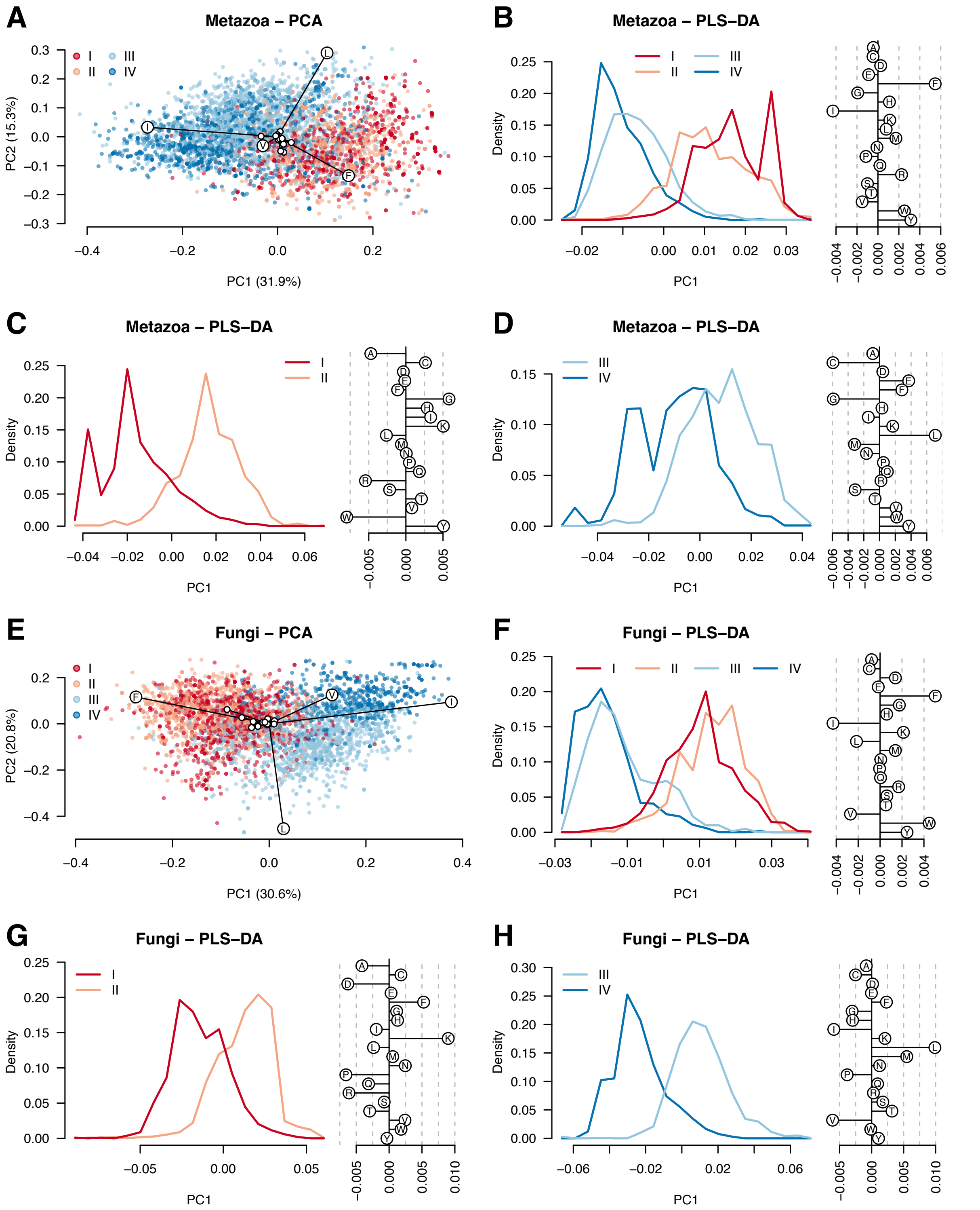


**Figure S3.** PCA and PLS-DA of Metazoan and fungi SNARE TMD amino acid composition. **(A)** PCA of SNARE TMD amino acid composition for Metazoa with corresponding loadings. Data points represent individual TMDs: I (1035 sequences), II (997), III (1869), IV (1602). **(B–D)** PLS-DA histograms of Metazoa TMDs. (B) Separation of classes I/II and III/IV. (C) Separation of classes I and II. The central peaks contain an excess of Qa SNAREs (Syx18 for I and Syx5 for II). The minor class I peak contains almost exclusively the R SNARE Sec22. (D) Separation of classes III and IV. Feature weights for each comparison are shown to the right. **(E)** PCA of SNARE TMD amino acid composition for fungi with corresponding loadings: I (1034 sequences), II (1137), III (1427), IV (665). **(F–H)** PLS-DA histograms of fungal TMDs: (F) I/II and III/IV; (G) I and II; (H) III and IV, with feature weights shown to the right.

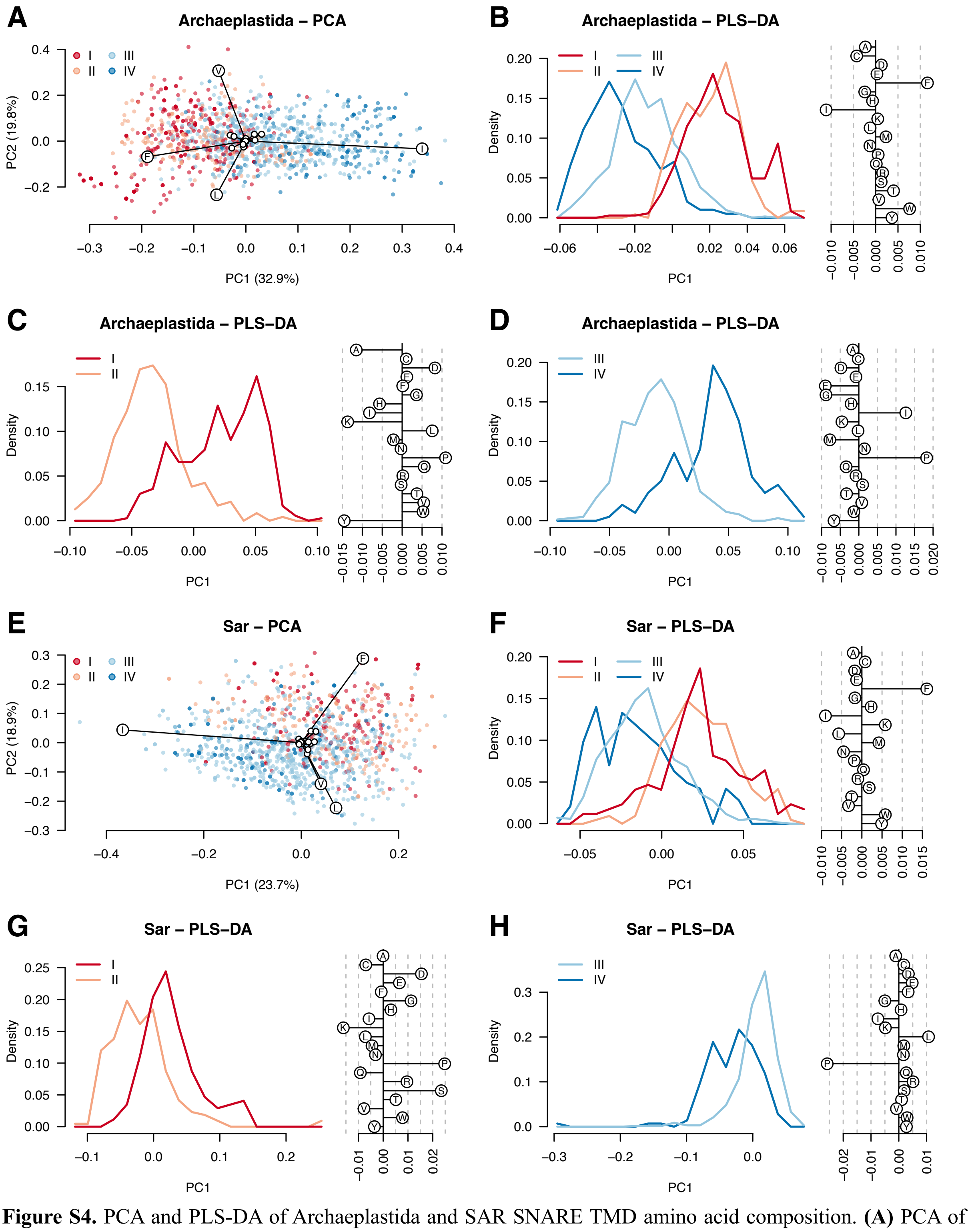


**Figure S4.** PCA and PLS-DA of Archaeplastida and SAR SNARE TMD amino acid composition. **(A)** PCA of SNARE TMD amino acid composition for Archaeplastida with corresponding loadings. Each point represents one TMD: I (365 sequences), II (236), III (622), IV (199). **(B–D)** PLS-DA histograms of Archaeplastida TMDs. (B) Separation of classes I/II and III/IV. (C) Separation of classes I and II. (D) Separation of classes III and IV. Feature weights for each comparison are shown to the right. **(E)** PCA of SAR SNARE TMD amino acid composition with corresponding loadings: I (172), II (217), III (690), IV (143). **(F–H)** PLS-DA histograms of SAR TMDs: (F) I/II and III/IV; (G) I and II; (H) III and IV, with feature weights shown to the right.

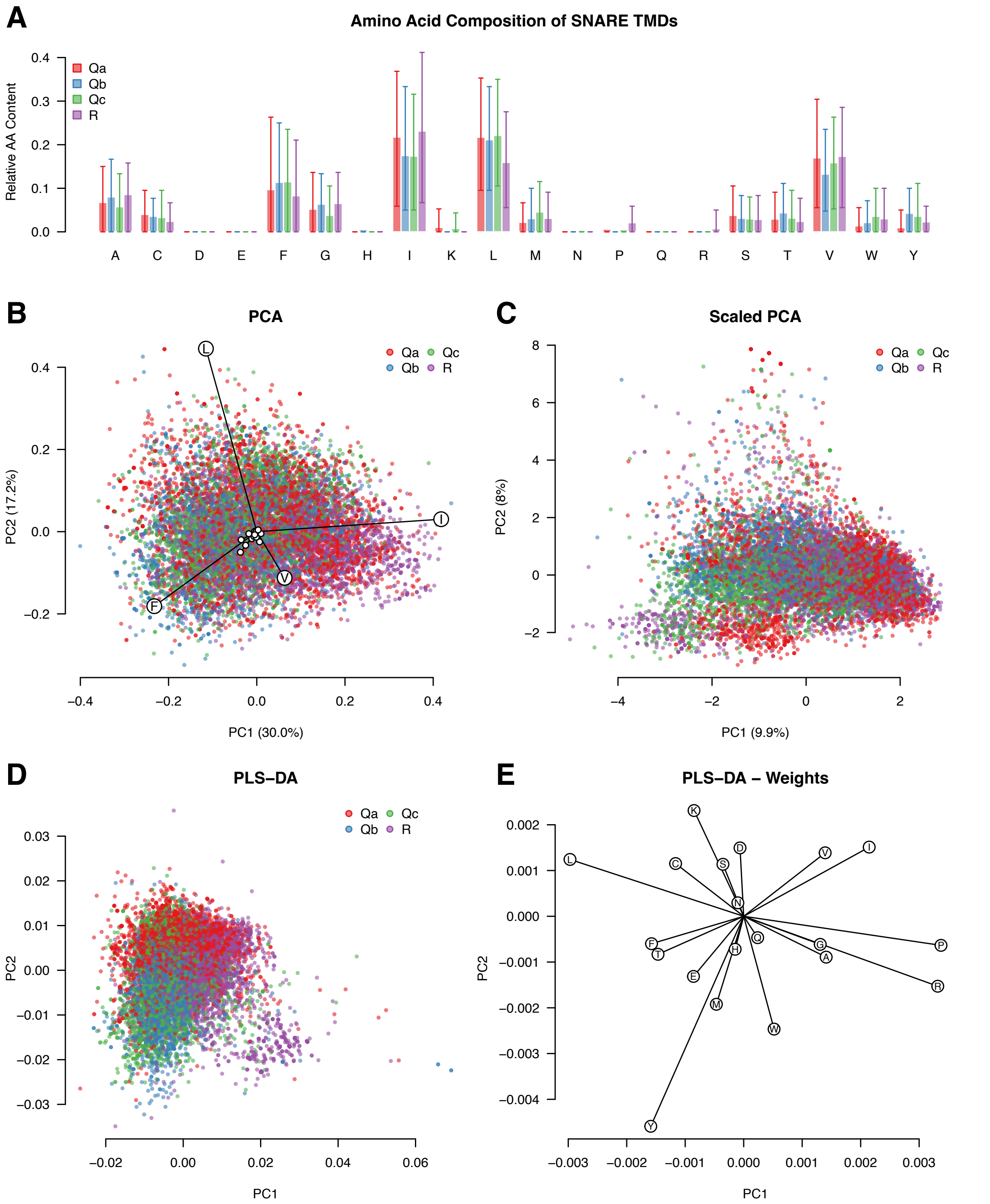


**Figure S5.** SNARE type-specific variation in TMD amino acid composition. **(A)** Mean amino acid composition of SNARE TMDs by SNARE type. Error bars indicate the range containing 80% of all TMDs. **(B)** Unscaled PCA of SNARE TMD amino acid composition with corresponding PCA loadings. Each point represents one TMD. **(C)** Scaled PCA of SNARE TMD amino acid composition. **(D)** Scaled PLS-DA separating all SNARE types using two principal components. **(E)** Feature weights contributing to the PLS-DA separation.

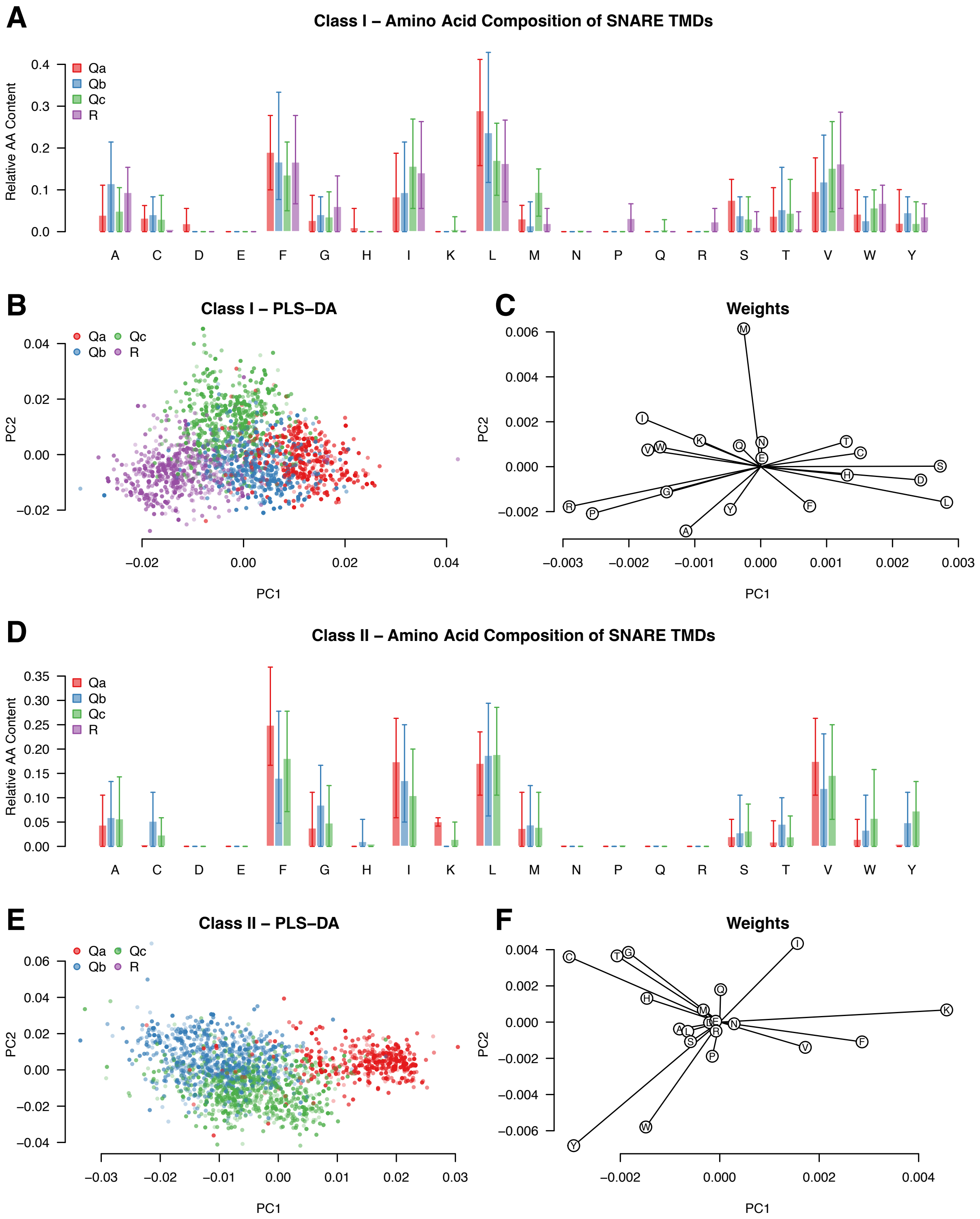


**Figure S6.** SNARE type-specific variation in TMD amino acid composition of Class I and Class II SNAREs. **(A)** Mean amino acid composition of class I SNARE TMDs. Error bars indicate the range containing 80% of all TMDs. **(B)** Scaled PLS-DA of class I SNARE TMD amino acid composition. Each point represents a single TMD. **(C)** Feature weights contributing to the class I PLS-DA separation. **(D)** Mean amino acid composition of class II SNARE TMDs. Error bars indicate the range containing 80% of all TMDs. **(E)** Scaled PLS-DA of class II SNARE TMD amino acid composition. Each point represents a single TMD. **(F)** Feature weights contributing to the class II PLS-DA separation. Data were resampled by drawing 2,000 sequences per class with replacement.

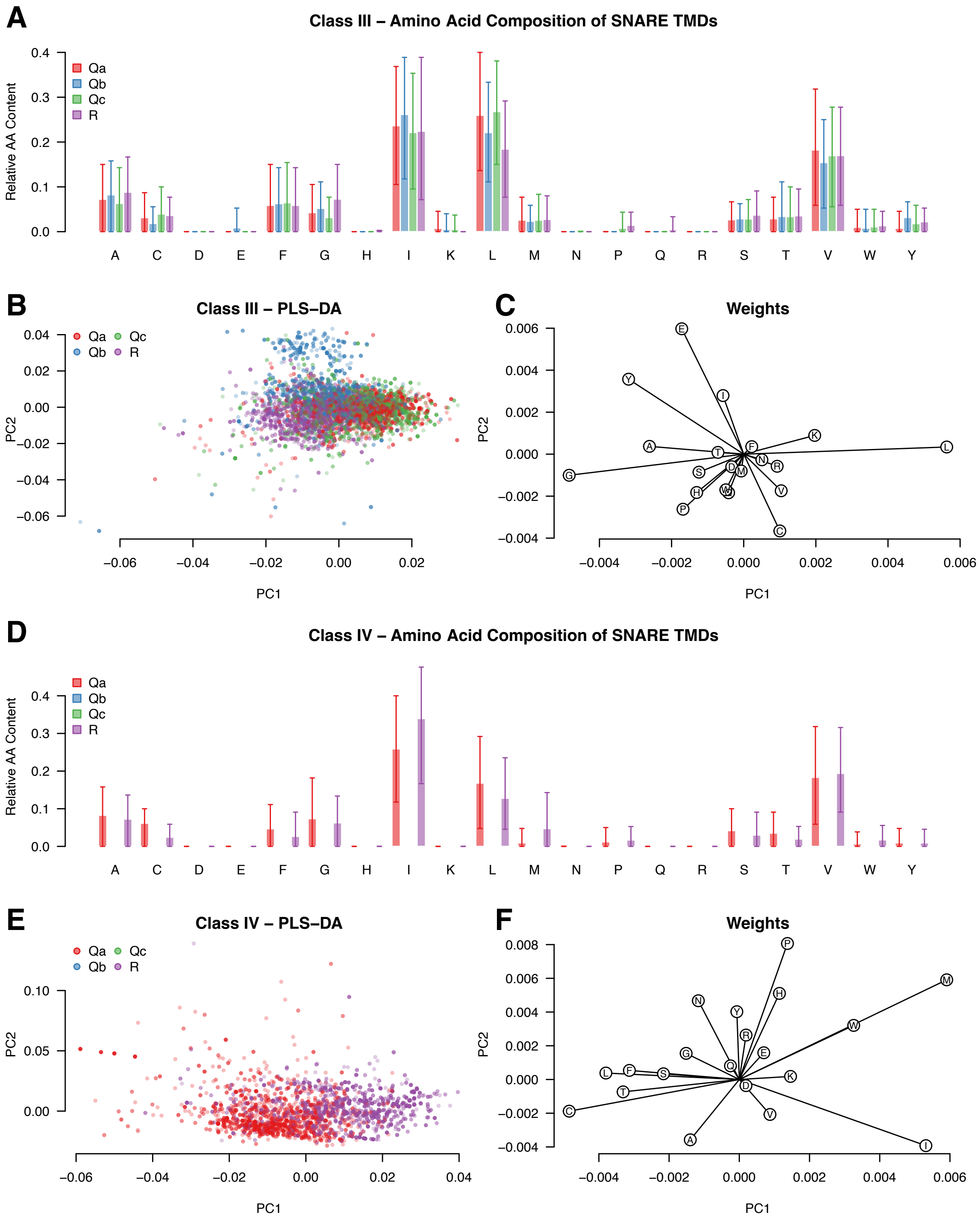


**Figure S7.** SNARE type-specific variation in TMD amino acid composition of Class III and Class IV SNAREs. **(A)** Mean amino acid composition of class III SNARE TMDs. Error bars indicate the range containing 80% of all TMDs. **(B)** Scaled PLS-DA of class III SNARE TMD amino acid composition. Each point represents a single TMD. **(C)** Feature weights contributing to the class III PLS-DA separation. **(D)** Mean amino acid composition of class IV SNARE TMDs. Error bars indicate the range containing 80% of all TMDs. **(E)** Scaled PLS-DA of class IV SNARE TMD amino acid composition. Each point represents a single TMD. **(F)** Feature weights contributing to the class IV PLS-DA separation. Data were resampled by drawing 2,000 sequences per class with replacement.

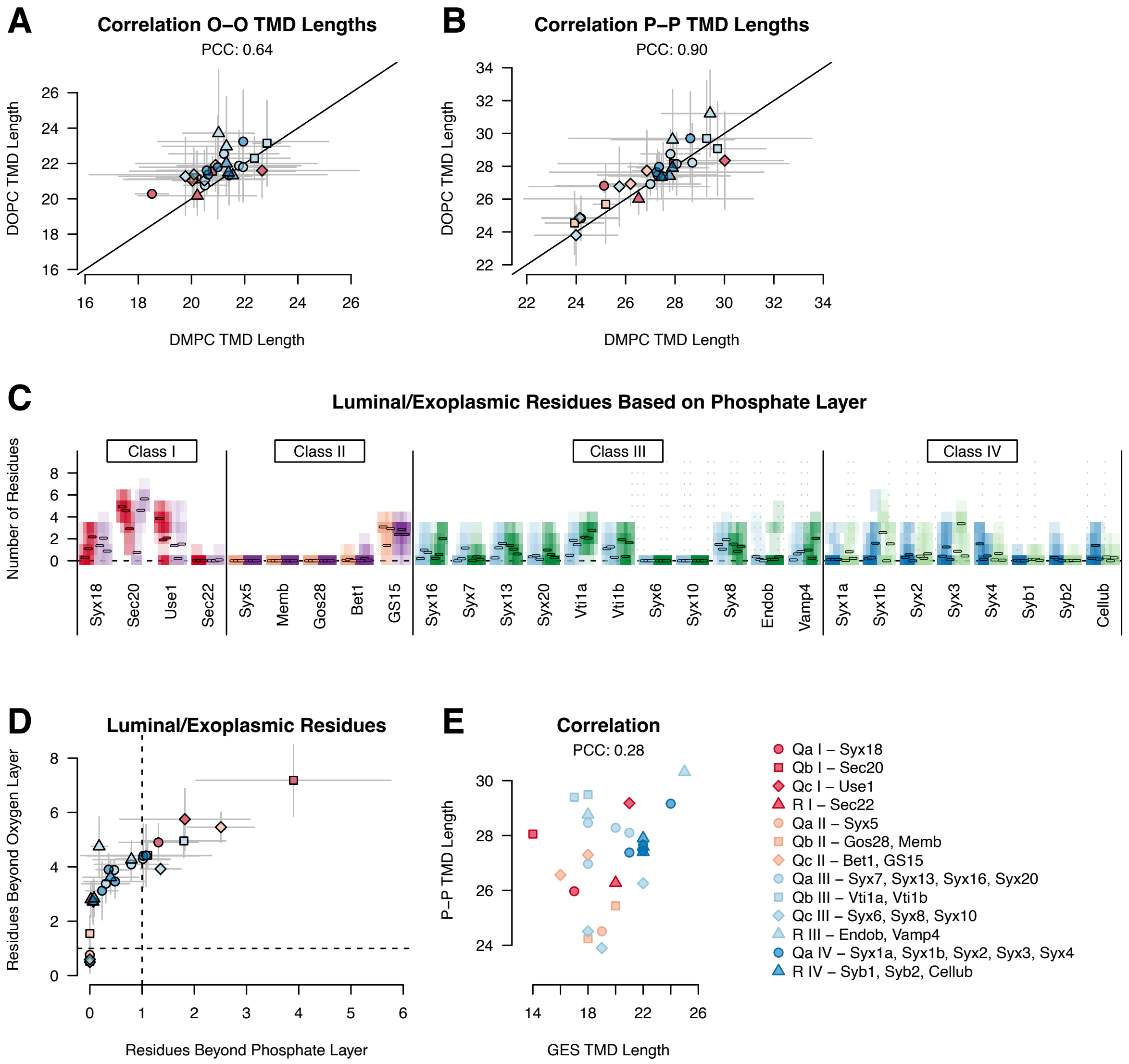


**Figure S8.** Comparison of TMD lengths from MD simulations and sequence-based predictions. **(A)** Correlation of O–O TMD lengths between DOPC and DMPC membranes. Data points indicate averages, and error bars show the 95% confidence intervals based on the three replicates per membrane. PCC is shown. **(B)** Correlation of P–P TMD lengths between DOPC and DMPC membranes. Averages, 95% confidence intervals, and PCC are shown. **(C)** Histogram of the number of *C*-terminal residues extending beyond the phosphate layer. Color intensity indicates frequency across trajectories; short bars denote averages per trajectory. **(D)** Average number of *C*-terminal residues outside the membrane using phosphate and oxygen layers. Error bars indicate the 95% confidence intervals. Dashed lines correspond to the length of a single residue. **(E)** Correlation between MD-derived P–P TMD lengths and sequence-based lengths obtained using the GES hydrophobicity scale.

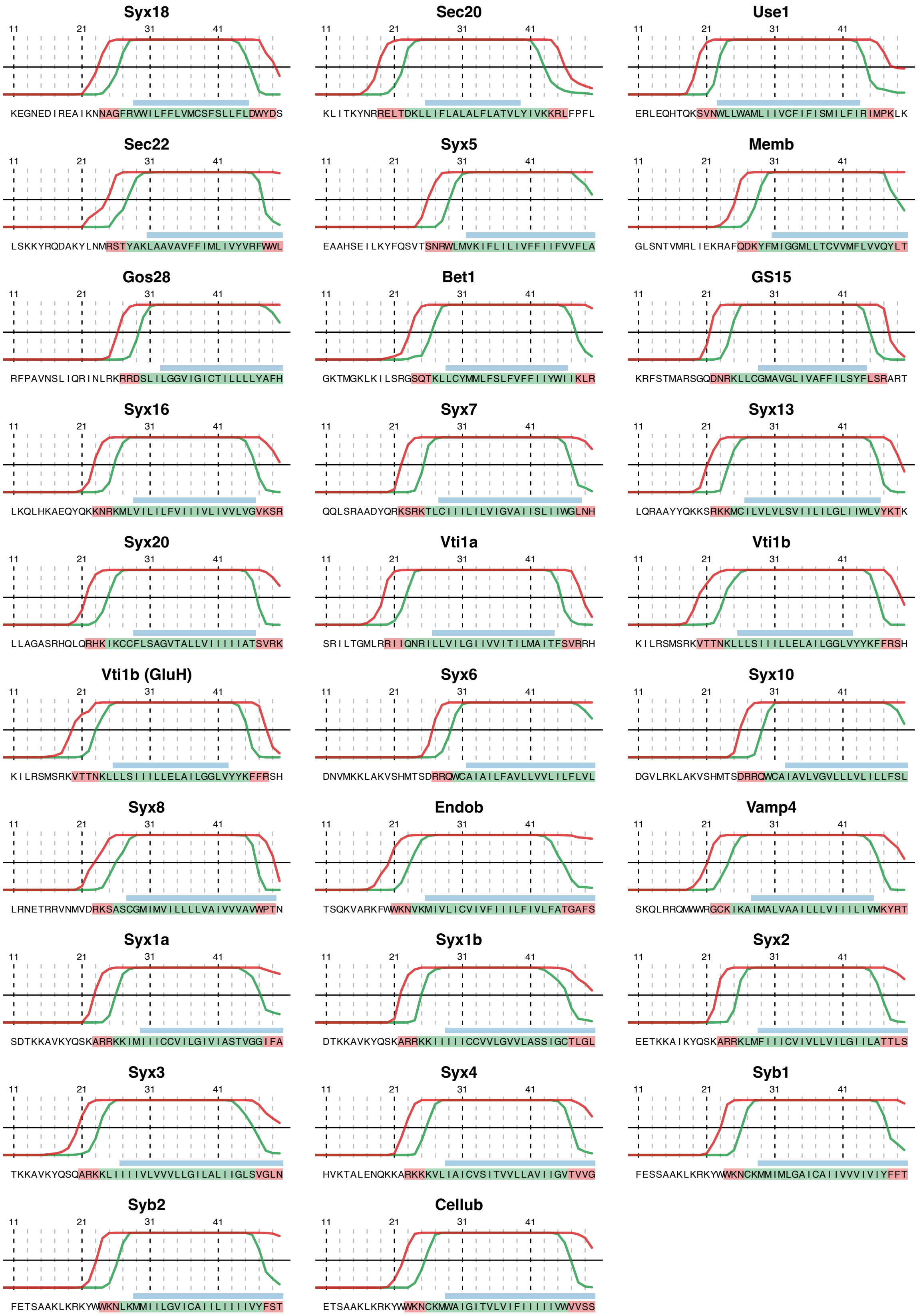


**Figure S9.** TMDs determined by MD simulations and the GES scale. The *C*-terminal 40 residues of each SNARE protein are shown. Curves indicate the fraction of frames in which each residue resides within the membrane across all trajectories, using the O–O definition (green) and P–P definition (red). Residues located within the membrane in ≥50% of frames are color-coded in the sequence accordingly (green for O–O, red for P–P). The blue bar highlights residues assigned to the TMD by the GES hydrophobicity scale. Vti1b is shown for both the charged and protonated (GluH) states.

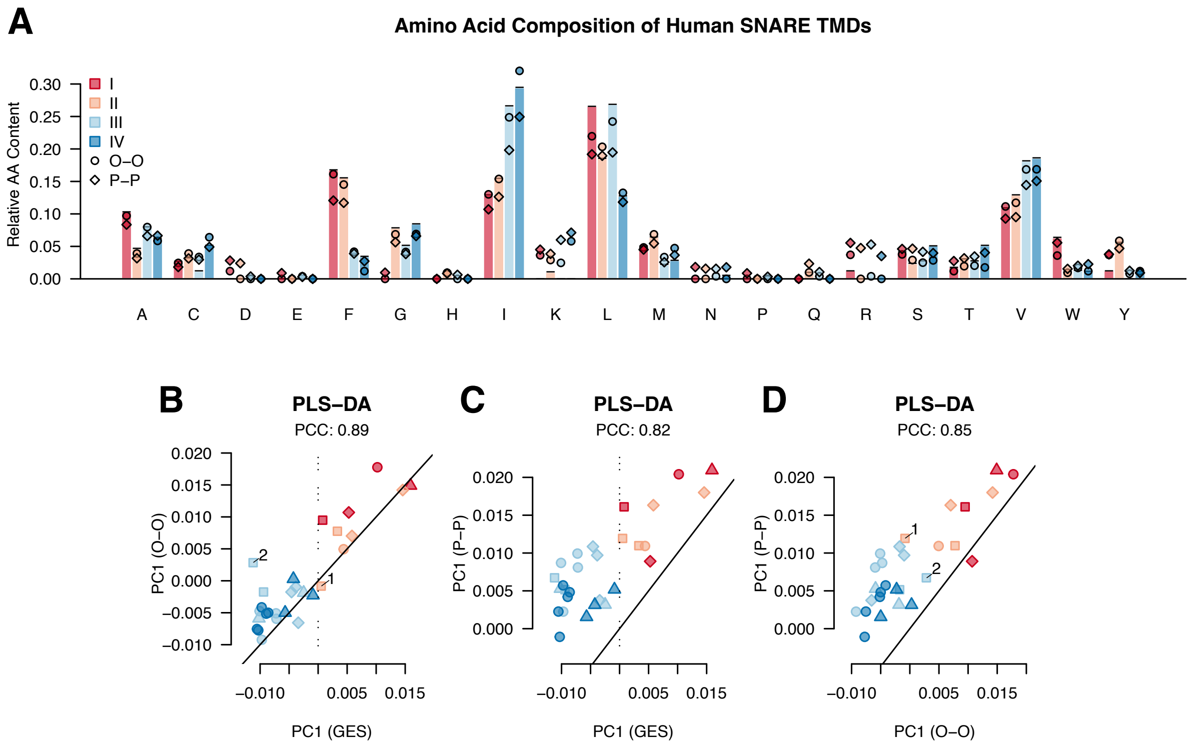


**Figure S10.** Amino acid composition of GES- and MD-defined SNARE TMDs. **(A)** Mean amino acid composition of human SNARE TMDs when determined using either the GES hydrophobicity scale and MD simulations. **(B)** Correlation of PLS-DA PC1 scores between TMDs defined by the O–O definition and the GES scale. PC1 scores were calculated using feature weights from the PLS-DA of the complete SNARE TMD dataset. Gos28 (1) and Vti1b (2) are highlighted. The dotted line highlights the separation between classes I/II and III/IV. **(C)** Correlation of PLS-DA PC1 scores between TMDs defined by the P–P definition and the GES scale. The dotted line highlights the separation between classes I/II and III/IV. **(D)** Correlation of PLS-DA PC1 scores between TMDs defined by the P–P and O–O definitions. Gos28 (1) and Vti1b (2) are highlighted.

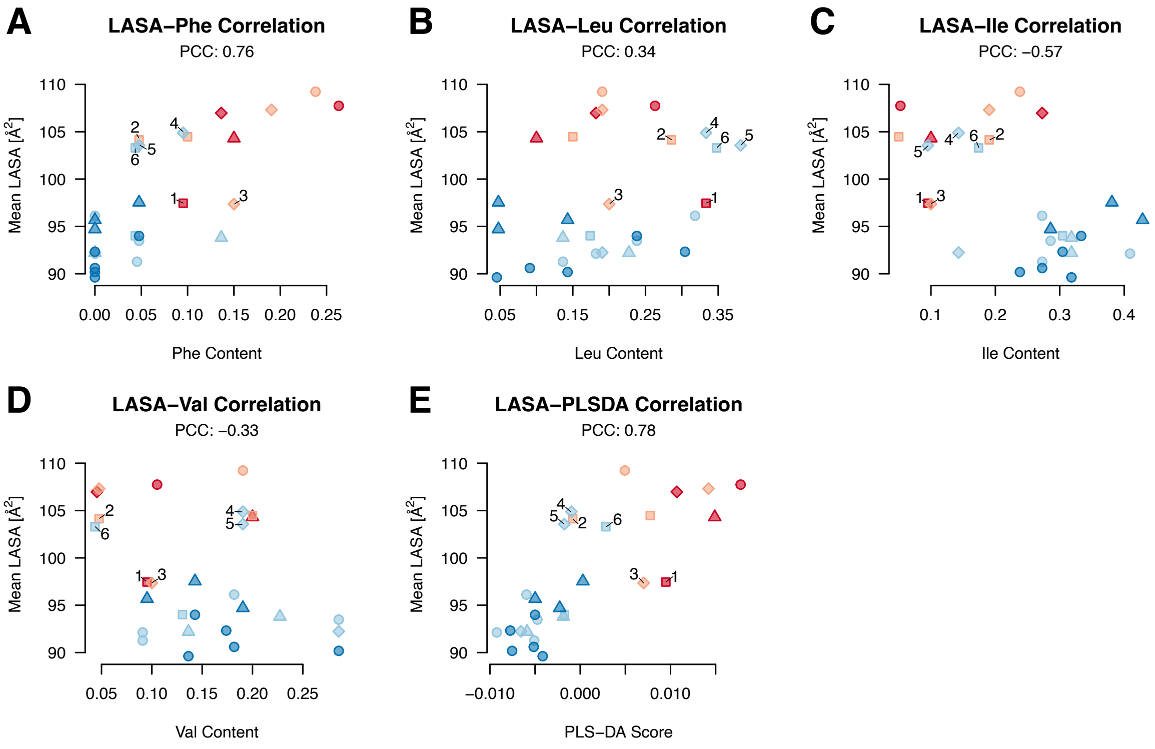


**Figure S11.** Correlations between LASA and amino acid composition in human SNARE TMDs based on the O–O definition: **(A)** LASA versus Phe content. **(B)** LASA versus Leu content. **(C)** LASA versus Ile content. **(D)** LASA versus Val content. **(E)** LASA versus PLS-DA scores. PCCs are indicated at the top of each plot. Highlighted SNAREs: Sec20 (1), Gos28 (2), GS15 (3), Syx6 (4), Syx10 (5), Vti1b (6).

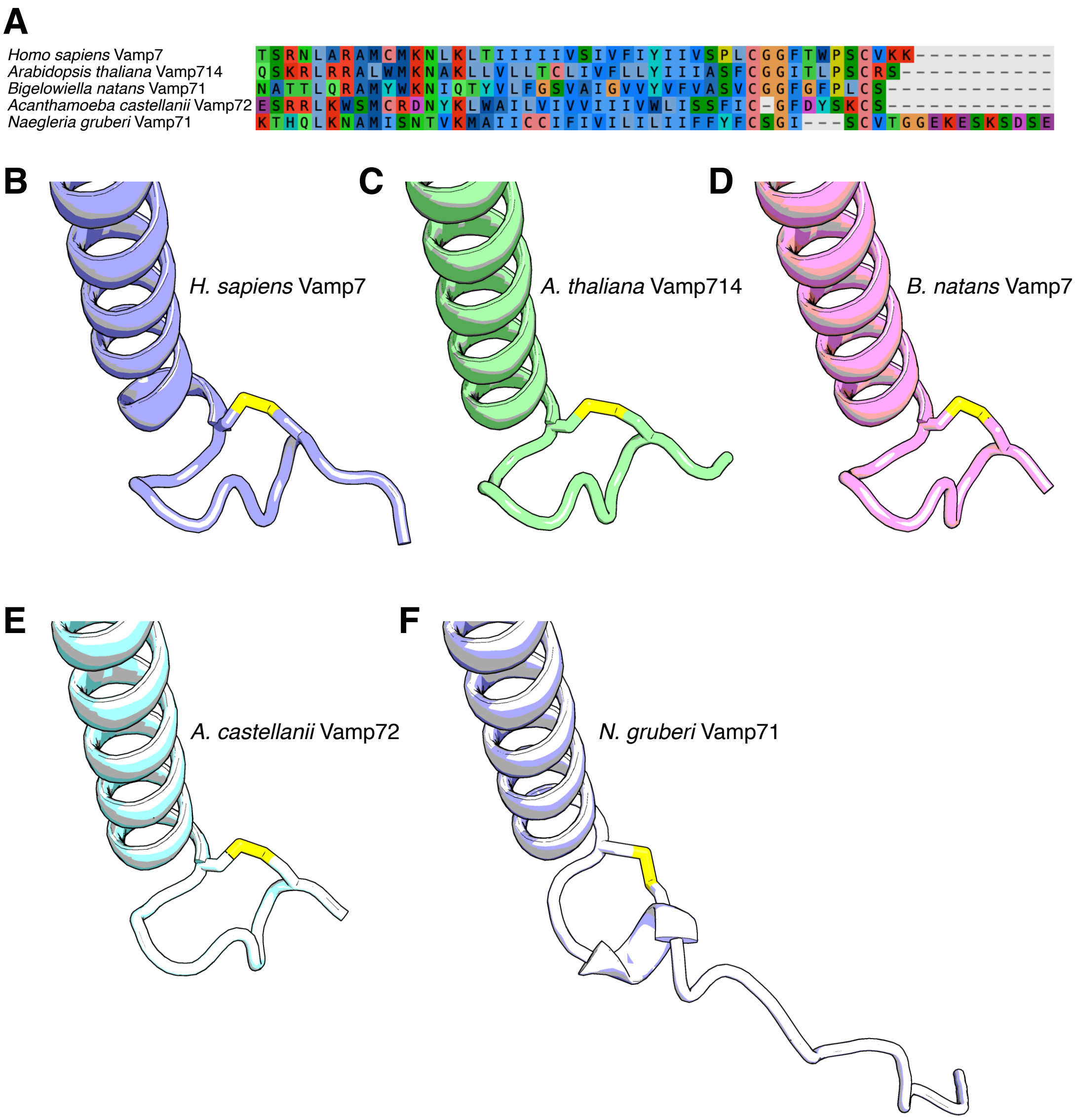


**Figure S12.** Conservation of the *C*-terminal region of Vamp7 across eukaryotes. **(A)** Multiple sequence alignment of the *C*-terminal regions of Vamp7 orthologs across Eukaryota. AF2 structural models are shown for **(B)** *Homo sapiens* (Obazoa) Vamp7, **(C)** *Arabidopsis thaliana* (Archaeplastida) Vamp714, **(D)** *Bigelowiella natans* (SAR) Vamp71, **(E)** *Acanthamoeba castellanii* (Amoebozoa) Vamp72, and **(F)** *Naegleria gruberi* (Discoba) Vamp71. Sulfur atoms forming the potentially conserved disulfide bond are highlighted in yellow.

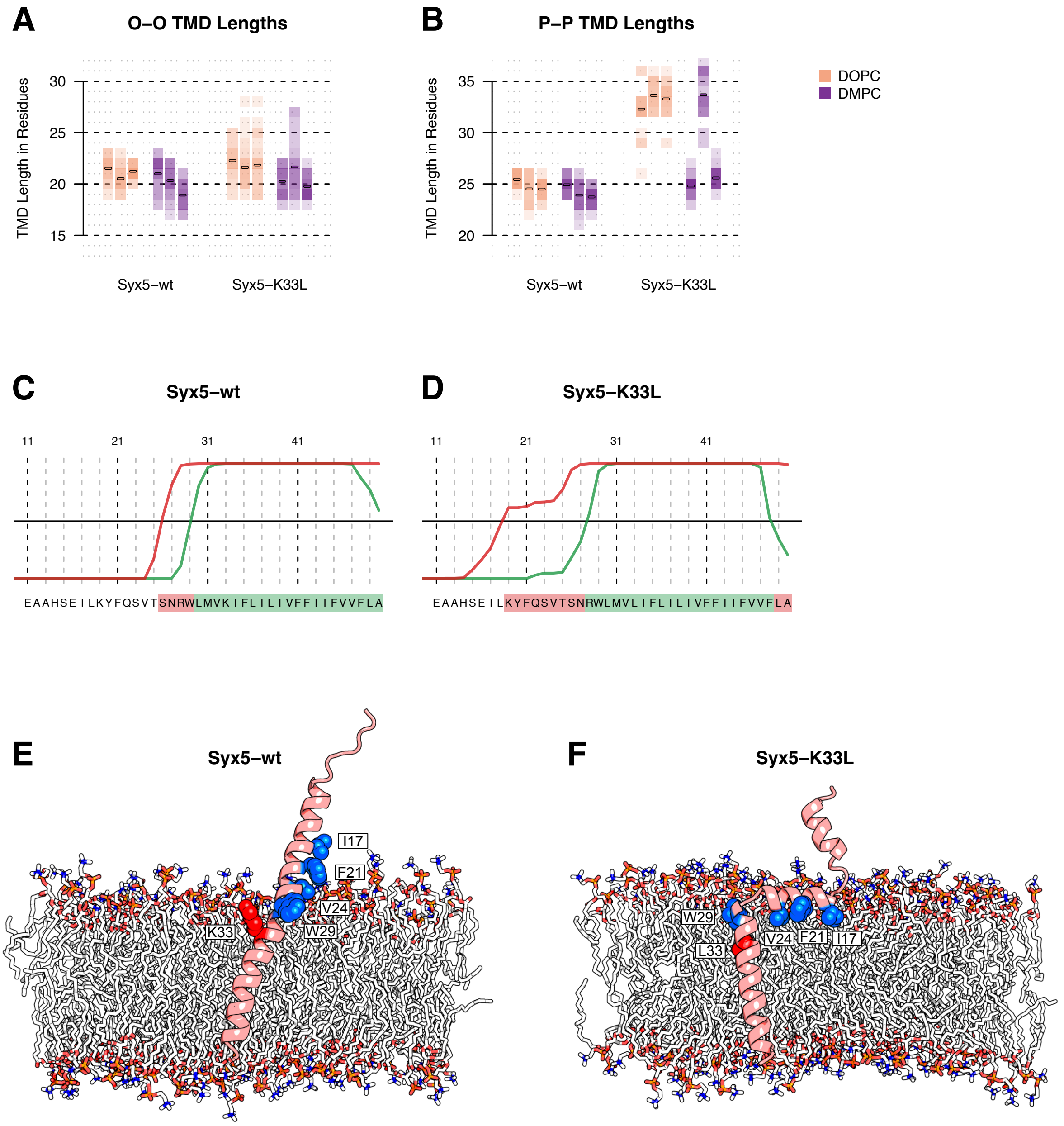


**Figure S13.** Effect of the K33L substitution on Syx5 TMD length and membrane insertion. **(A)** Histogram of TMD lengths based on the P–P definition for wt- and K33L-Syx5 trajectories. Color intensity indicates how frequently a given TMD length was observed. Small bars indicate the mean for each trajectory. **(B)** Same as (A), but using the O–O definition. Most K33L-Syx5 trajectories display an increase in TMD length due to more residues being inserted on the cytosolic side of the membrane. **(C, D)** TMDs determined by MD simulations. The *C*-terminal 40 residues of wt-Syx5 (C) and K33L-Syx5 (D) are shown. Curves indicate the fraction of frames in which each residue resides within the membrane across all trajectories, using the O–O (green) and P–P definition (red). Residues located within the membrane in ≥50% of frames are color-coded in the sequence accordingly (green for O–O, red for P–P). **(E, F)** MD snapshots of wt-Syx5 (E) and K33L-Syx5 (F) obtained from simulations with DOPC to illustrate the sampled conformational states. The interfacial Trp residue and the hydrophobic residues of the amphipathic helix are highlighted in blue. The Lys residue in the wt TMD and its Leu substitution are shown in red. The K33L substitution promotes deeper insertion of the TMD into the membrane and induces a pronounced kink within the helix, resulting in partial insertion of the TMD-adjacent region through the hydrophobic face of the amphipathic helix. These findings suggest that K33 contributes to proper positioning of the TMD within the membrane.

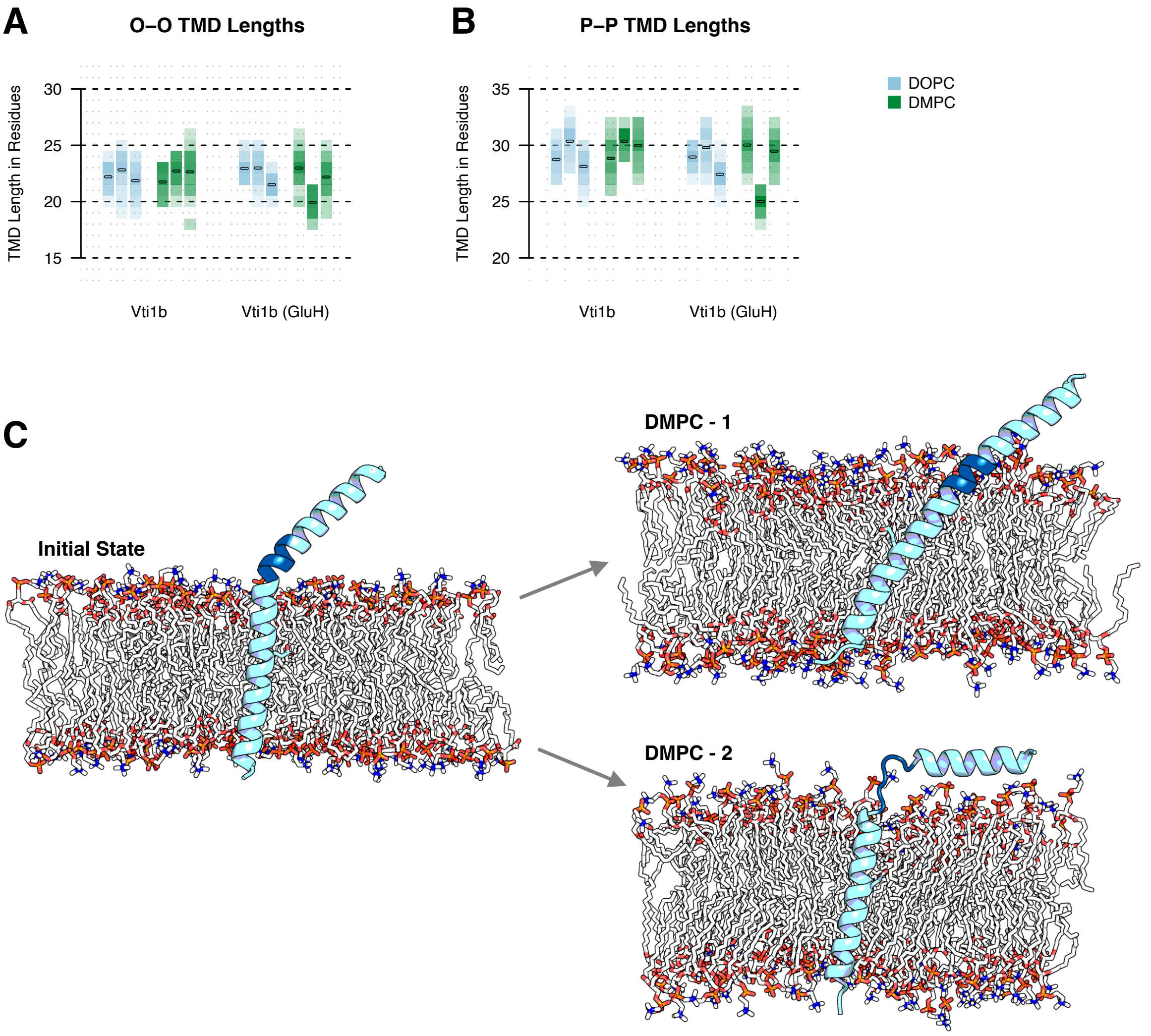


**Figure S14.** Influence of Glu protonation on Vti1b TMD length. **(A)** Histogram of TMD lengths based on the P–P definition for all Vti1b trajectories with Glu in its charged and in its neutral (GluH) state. Color intensity indicates how frequently a given TMD length was observed. Small bars indicate the mean for each trajectory. Differences between the two Glu protonation states were not statistically significant (two-tailed t-test, $p = 0.65$). **(B)** Same as (A), but using the O–O definition ($p = 0.31$). One Vti1b (GluH) trajectory in DMPC displays a markedly shorter TMD due to formation of a loop between the TMD and the *N*-terminus. **(C)** MD snapshots of Vti1b obtained from DMPC simulations illustrating the sampled conformation states. The loop observed in the second DMPC trajectory (residues 18–RKVTTN–23) is highlighted in dark blue. Similar rearrangements of the *N*-terminal region occurred in other SNARE simulations, such as the first trajectory of Syx1b in DMPC. Looped or kinked states have also been reported previously.[1]

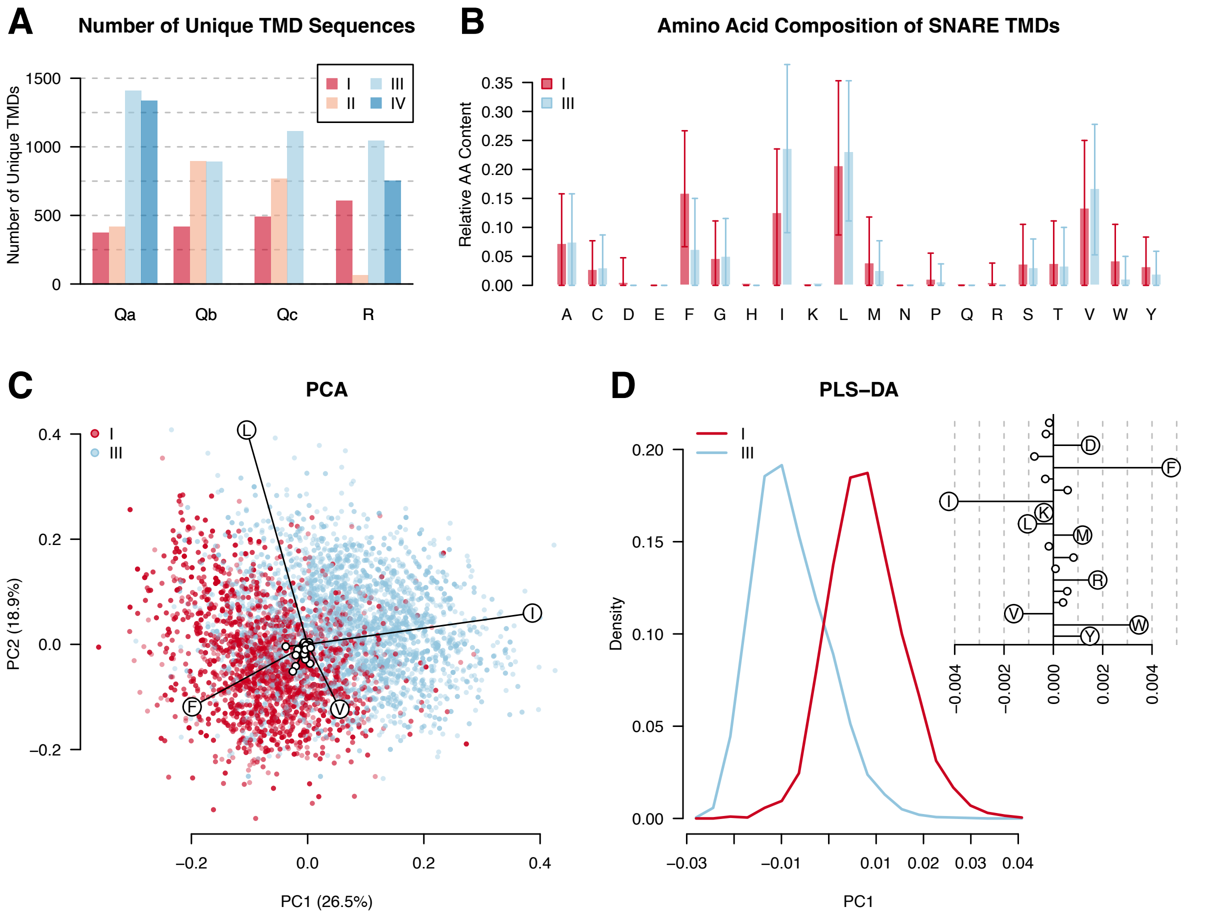


**Figure S15.** Subsampling analysis of SNARE TMD amino acid composition. **(A)** Number of unique SNARE TMD sequences grouped by type (Qa, Qb, Qc, R) and class (I–IV). Only class I and III contain all four SNARE types and were therefore used for this analysis. For each type-class combination (e.g., Qa II), 1,000 TMDs were randomly sampled with replacement. **(B)** Mean amino acid composition of class I and class III SNARE TMDs. TMDs were defined using the GES hydrophobicity scale. Error bars indicate the range containing 80% of all TMDs. **(C)** PCA of SNARE TMD amino acid composition with corresponding relative loadings. Each point represents one TMD sequence. **(D)** Histogram of PLS-DA results separating classes I/II and III/IV, with feature weights shown in the upper-right panel.

**Alignment S1.** *C*-terminal sequences of human SNAREs used for MD simulations in this study.

```
>01_Syx18
QLVVGATENIKEGNEDIREAIKNNAGFRVWILFFLVMCSFSLLFLDWYDS
>02_Sec20
SMSGTIQLGRKLITKYNRRELTDKLLIFLALALFLATVLYIVKKRLFPFL
>03_Use1
QNLEKLKTESERLEQHTQKSVNWLLWAMLIIVCFIFISMILFIRIMPKLK
>04_Sec22
LDSKANNLSSLSKKYRQDAKYLNMRSTYAKLAAVAVFFIMLIVYVRFWWL
>05_Syx5
ENVLGAQLDVEAAHSEILKYFQSVTSNRWLMVKIFLILIVFFIIFVVFLA
>06_Memb
KKILDIANMLGLSNTVMRLIEKRAFQDKYFMIGGMLLTCVVMFLVVQYLT
>07_Gos28
IHSKMNTLANRFPAVNSLIQRINLRKRRDSLILGGVIGICTILLLLYAFH
>08_Bet1
SQFDSTTGFLGKTMGKLKILSRGSQTKLLCYMMLFSLFVFFIIYWIIKLR
>09_GS15
SMTSLLTGSVKRFSTMARSGQDNRKLLCGMAVGLIVAFFILSYFLSRART
>10_Syx16
EQSCIKTEDGLKQLHKAEQYQKKNRKMLVILILFVIIIVLIVVLVGVKSR
>11_Syx7
NAEVHVQQANQQLSRAADYQRKSRKTLCIIILILVIGVAIISLIIWGLNH
>12_Syx13
EVHVERATEQLQRAAYYQKKSRKKMCILVLVLSVIILILGLIIWLVYKTK
>13_Syx20
ASSHAEAARQLLAGASRHQLQRHKIKCCFLSAGVTALLVIIIIIATSVRK
>14_Vti1a
RETDANLGKSSRILTGMLRRIIQNRILLVILGIIVVITILMAITFSVRRH
>15_Vti1b
NTSENLSKSRKILRSMSRKVTTNKLLLSIIILLELAILGGLVYYKFFRSH
>16_Syx6
HELESTQSRLDNVMKKLAKVSHMTSDRRQWCAIAILFAVLLVVLILFLVL
>17_Syx10
QEMDHTQSRMDGVLRKLAKVSHMTSDRRQWCAIAVLVGVLLLVLILLFSL
>18_Syx8
ANLVENTDEKLRNETRRVNMVDRKSASCGMIMVILLLLVAIVVVAVWPTN
>19_Endob
LEATSEHFKTTSQKVARKFWWKNVKMIVLICVIVFIIILFIVLFATGAFS
>20_Vamp4
SDNATAFSNRSKQLRRQMWWRGCKIKAIMALVAAILLLVIIILIVMKYRT
>21_Vamp7
FKTTSRNLARAMCMKNLKLTIIIIIVSIVFIYIIVSPLCGGFTWPSCVKK
>22_Syx1a
HAVDYVERAVSDTKKAVKYQSKARRKKIMIIICCVILGIVIASTVGGIFA
>23_Syx1b
SVDYVERAVSDTKKAVKYQSKARRKKIIIIICCVVLGVVLASSIGCTLGL
>24_Syx2
NATDYVEHAKEETKKAIKYQSKARRKLMFIIICVIVLLVILGIILATTLS
>25_Syx3
VDHVEKARDETKKAVKYQSQARKKLIIIIVLVVVLLGILALIIGLSVGLN
>26_Syx4
SADYVERGQEHVKTALENQKKARKKKVLIAICVSITVVLLAVIIGVTVVG
>27_Syb1
ADALQAGASQFESSAAKLKRKYWWKNCKMMIMLGAICAIIVVVIVIYFFT
>28_Syb2
ADALQAGASQFETSAAKLKRKYWWKNLKMMIILGVICAIILIIIIVYFST
>29_Cellub
DALQAGASQFETSAAKLKRKYWWKNCKMWAIGITVLVIFIIIIIVWVVSS
>30_Myob
MSSTFNKTTQNLAQKKCWENIRYRICVGLVVVGVLLIILIVLLVVFLPQSSDSSS
```
